# Logging alters tropical forest structure, while conversion to agriculture reduces biodiversity and functioning

**DOI:** 10.1101/2022.12.15.520573

**Authors:** Charles J. Marsh, Edgar C. Turner, Benjamin Wong Blonder, Boris Bongalov, Sabine Both, Rudi S. Cruz, Dafydd M. O. Elias, David Hemprich-Bennett, Palasiah Jotan, Victoria Kemp, Ully H. Kritzler, Sol Milne, David T. Milodowski, Simon L. Mitchell, Milenka Montoya Pillco, Matheus Henrique Nunes, Terhi Riutta, Samuel J. B. Robinson, Eleanor M. Slade, Henry Bernard, David F. R. P. Burslem, Arthur Y. C. Chung, Elizabeth L. Clare, David A. Coomes, Zoe G. Davies, David P. Edwards, David Johnson, Pavel Kratina, Yadvinder Malhi, Noreen Majalap, Reuben Nilus, Nicholas J. Ostle, Stephen J. Rossiter, Matthew J. Struebig, Joseph A. Tobias, Mathew Williams, Robert M. Ewers, Owen T. Lewis, Glen Reynolds, Yit Arn Teh, Andy Hector

**Affiliations:** Department of Biology, University of Oxford; Oxford, UK; University Museum of Zoology, University of Cambridge; Cambridge, UK; Department of Environmental Science, Policy, and Management, University of California Berkeley; Berkeley, CA, USA; Department of Plant Sciences and Conservation Research Institute, University of Cambridge; Cambridge, UK; School of Environmental and Rural Science, University of New England; Armidale, Australia; Escuela de Biología, Universidad Nacional San Antonio Abad del Cusco; Cuzco, Peru; UK Centre for Ecology & Hydrology; Lancaster Environment Centre, Lancaster, UK; School of Biological and Behavioural Sciences, Queen Mary University of London; London, UK; Department of Forest Ecology, Faculty of Forestry and Wood Sciences, Czech University of Life Sciences Prague; Czech Republic; Department of Earth and Environmental Sciences, University of Manchester; Manchester, UK; School of Biological Sciences, University of Aberdeen; Aberdeen, UK; School of GeoSciences and NCEO, University of Edinburgh; Edinburgh, UK; Durrell Institute for Conservation & Ecology, School of Anthropology and Conservation, University of Kent; Canterbury, UK; Institute of Biological and Environmental Sciences, University of Aberdeen; Aberdeen, UK; Department of Geographical Sciences, University of Maryland, College Park, MD, USA; Environmental Change Institute & Leverhulme Centre for Nature Recovery, School of Geography and the Environment, University of Oxford; Oxford, UK; UK Centre for Ecology and Hydrology, Wallingford, UK; Faculty of Natural Sciences, Imperial College; London, UK; Lancaster Environment Centre, Lancaster University; Lancaster, UK; Asian School of the Environment, Nanyang Technological University; Singapore; Institute for Tropical Biology and Conservation, Universiti Malaysia Sabah; Kota Kinabalu, Sabah, Malaysia; Forest Research Centre, Sabah Forestry Department; Sandakan, Sabah, Malaysia; Department of Biology, York University; Toronto, ON, Canada; Centre for Global Wood Security, University of Cambridge, Cambridge, UK; Southeast Asia Rainforest Research Partnership; Lahad Datu, Sabah, Malaysia; School of Natural and Environmental Sciences, Newcastle University; Newcastle upon Tyne, UK; Department of Biology & Leverhulme Centre for Nature Recovery, University of Oxford; Oxford, UK

## Abstract

The impacts of degradation and deforestation on tropical forests are poorly understood, particularly at landscape scales. We present an extensive ecosystem analysis of the impacts of logging and conversion of tropical forest to oil palm from a large-scale study in Borneo, synthesizing responses from 82 variables categorized into four ecological levels spanning a broad suite of ecosystem properties: 1) structure and environment, 2) species traits, 3) biodiversity, and 4) ecosystem functions. Responses were highly heterogeneous and often complex and non-linear. Variables that were directly impacted by the physical process of timber extraction, such as soil structure, were sensitive to even moderate amounts of logging, whereas measures of biodiversity and ecosystem functioning were generally resilient to logging but more affected by conversion to oil palm plantation.

**One-Sentence Summary:** Logging tropical forest mostly impacts structure while biodiversity and functions are more vulnerable to habitat conversion.

---

Tropical forests support biodiversity and provide ecosystem services such as stocks and flows of carbon, nutrients and water, but their structure and functioning are threatened by degradation and conversion to other land uses (*1*, *2*). A major cause of tropical forest degradation is selective logging for timber which can increase vulnerability to subsequent deforestation (*3–5*). In Southeast Asia, many forests have experienced multiple rounds of selective logging, with some then converted to oil palm plantations (*6*, *7*), resulting in large-scale forest losses (3.25 Mha in Malaysia and Indonesia between 2000 and 2011 (*8*)) and increased carbon emissions (4,051 MtCO_2_ in the same countries over the same period (*8*)). Indeed, ∼45% of Southeast Asian oil palm plantations have been established through direct clearing of forest (*9*).

Knowledge of the full environmental impacts of logging and forest conversion in the tropics to other land uses such as oil palm (the forest disturbance gradient) is limited (*10–12*). The logistical challenges of studying highly biodiverse tropical forest ecosystems means that there are few comprehensive assessments of the impacts on biodiversity and the multiple ecosystem functions and services that tropical forests provide across the full disturbance gradient at the landscape scale (*13*). Here, we undertake a comprehensive assessment of how biodiversity, structure, and functioning of tropical forest ecosystems are altered across a disturbance gradient of increasing intensity of selective logging and conversion to oil palm plantation, and examine the points along that gradient where changes from old-growth forest conditions are most apparent.

We synthesize data from 82 metrics of ecosystem properties that collectively provide a comprehensive assessment of environmental and ecological conditions, capturing aspects of the forest structure and environment, as well as measures of biodiversity and ecosystem functioning. Data were collected as part of a coordinated large-scale study in the Stability of Altered Forest Ecosystems (SAFE) Project (*14*) and associated sites of the Human Modified Tropical Forests (HMTF) programme in the Malaysian state of Sabah, Borneo (Fig. 1a), where patterns of deforestation are representative of other regions in Southeast Asia (*15*). Study sites were located in areas of intact and disturbed lowland dipterocarp rainforest and oil palm plantations.

**Fig. 1.**
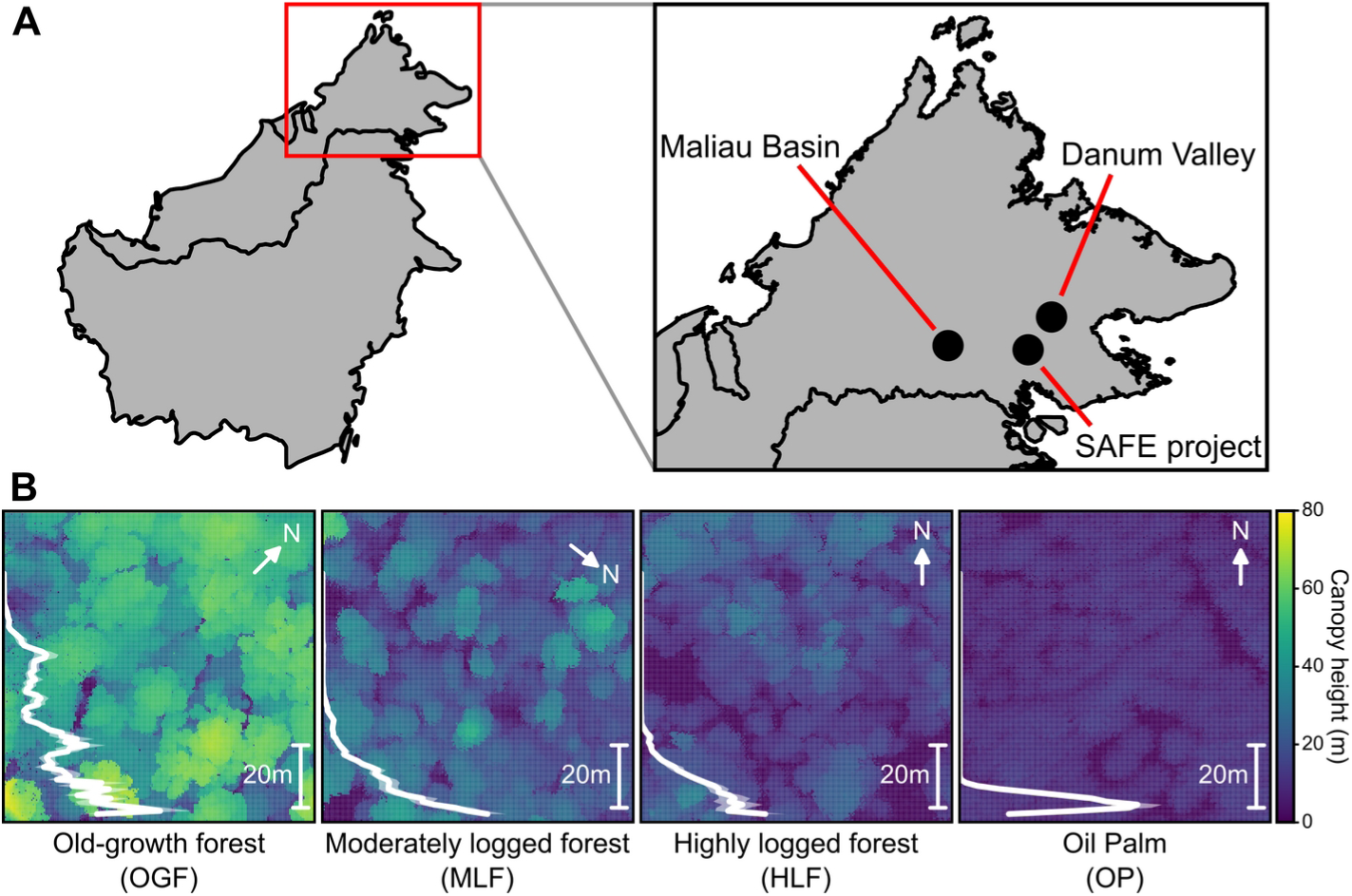
Study sites and disturbance categories. (**A**) Location of the study sites in Sabah, Malaysian Borneo. (**B**) Canopy height profiles of the study systems for representative 1 ha plots (from left to right): old-growth forest (OGF), moderately logged forest (MLF), highly logged forest (HLF), and oil palm plantation (OP). Backgrounds show the maximum canopy height for each pixel, and inset graphs show the plant area density (mean ± 95% C.I.) of the vertical forest structure estimated through LiDAR.

We use a replicated experimental design and standardized analyses (sample sizes ranging from 27 to 373,968 across the 82 variables; Table S1) to quantify the impacts of selective logging and land-use change across different intensities of disturbance from: (1) old-growth forest (OGF), through (2) moderately logged (MLF) and (3) highly logged (HLF) forest, to (4) oil palm plantation (OP). To allow us to synthesize the effects of habitat change on the whole ecosystem, we focus on understanding the comprehensive impacts of changes, rather than assessing specific drivers affecting each metric. Logged forest sites had an average of ca. 113 m^3^ ha^-1^ of timber harvested during 1978, with a second cycle of harvesting in the late 1990s to early 2000s, removing a further ∼66 m^3^ ha^-1^ in three rounds (HLF sites) or ∼37 m^3^ ha^-1^ in two rounds (MLF) (*16*). Forests along the disturbance intensity gradient were characterized by a decrease in basal area of mature trees, a more open canopy, fewer large trees, and a higher proportion of pioneer tree species (*17*). Measurements with airborne LiDAR showed a progressive reduction of canopy height and simplification of canopy structure from OGF to MLF and HLF (Fig. 1), culminating in a homogeneous, single low layer in oil palm (*18*).

## CATEGORIZING VARIABLES INTO ECOLOGICAL LEVELS

The 82 response variables (Tables S1-5) detail ecosystem properties sampled in OGF and one or more of the disturbed habitat categories. Each property was categorized into one of four ecological levels, building in complexity and distance from the direct impacts of logging (*17*). Although the assignment of responses to levels is partly subjective, they provide a useful framework for summarizing the ecosystem effects of logging and conversion, as each higher level generally captures features of properties at lower levels. Level 1 (Structure and Environment) comprised variables related to soil properties, microclimate, and forest structure that are directly affected by the physical processes of timber extraction and oil palm cultivation. Level 2 (Tree traits) constituted the traits of the remaining tree community, reflecting the change in plant species composition caused by selective logging, as well as subsequent colonization and growth of early successional species. Tree traits were grouped according to whether they contributed to structural stability and defense (structural traits), leaf photosynthetic potential and leaf longevity (photosynthesis traits), or foliar concentrations of key mineral nutrients (nutrient traits). Level 3 (Biodiversity) quantified below-and above-ground multi-trophic and functional biodiversity, from assemblages of soil microorganisms to consumers in higher trophic levels, that strongly depend on the abiotic and structural conditions described in level 1, and the tree community diversity and composition in level 2. Level 4 (Functioning) corresponded to ecosystem functions, such as decomposition, which, within a given environment, are largely defined by the composition of communities described in levels 2 and 3.

To allow comparison of multiple responses on a common scale, we transformed the raw data if necessary to improve normality and then standardized all variables as z-scores (mean-centering each variable by subtracting its mean value and dividing by its standard deviation) before analyzing them using linear mixed-effects models to assess effects across the disturbance gradient (the four disturbance categories: OGF, MLF, HLF and OP), while taking account of the spatial hierarchical structure of our datasets (Fig. S1). To provide a comprehensive assessment, where possible, datasets were analyzed across multiple facets and spatial scales (Table S1). For example, we calculated three measures of the effective number of species for some biodiversity datasets (*19*): effort-standardized species richness (Hill number *q* = 0), Shannon diversity (*q* = 1), and Simpson’s diversity (*q* = 2). Similarly, we analyzed the species richness of some groups at the finest spatial grain at which those data were collected, but also aggregated data to coarser grain sizes. Although positive or negative responses for some variables have clear desirable or undesirable consequences, for others such as β-diversity, there is no obvious value judgment. We therefore focus on examining whether each variable changed from values recorded within OGF, and where along the gradient change occurred.

## STRONG BUT HETEROGENEOUS RESPONSES TO DISTURBANCE ACROSS ECOLOGICAL LEVELS

Overall, 60 of the 82 response variables showed statistically significant differences across the disturbance intensity gradient (Likelihood Ratio Tests (*LRT*) against a null model with no disturbance factor; Table S6). This was far greater than the expected level of false positives (∼4 out of 82 with a significance level of *p* < 0.05) in the absence of any effect of disturbance (Fig. 2, solid lines).This result was consistent when controlling for dataset identity through randomization (84.27 % of variables significant) (*17*). The proportion of significant results and the degree of variation explained by disturbance intensity (the mean marginal *R*^2^ from linear mixed-effects models for group; Table S6), varied with the ecological level of the response variable. Generally, the responses that showed the strongest effect of disturbance were those most directly affected by logging (*18*) — level 1, environment and structure (mean marginal *R*^2^ ± s.e. for datasets with OP included = 0.210 ± 0.034, 9 out of 12 variables LRT *p* < 0.05; OP excluded = 0.228 ± 0.068, all 4 variables LRT *p* < 0.05) and level 2, aggregated tree traits (first axis values from a PCA; OP excluded = 0.253 ± 0.029, all 3 variables LRT *p* < 0.05). Biodiversity measures (level 3) showed stronger responses to disturbance intensity in variables where oil palm was sampled than those measured in forest habitats only (OP included = 0.232 ± 0.023, 11 out of 13 variables LRT *p* < 0.05; OP excluded = 0.113 ± 0.021, 11 out of 12 variables LRT *p* < 0.05). Ecosystem functioning variables (level 4) showed weaker responses to disturbance intensity overall, with little change across the gradient for some variables (OP included = 0.081 ±0.021, 2 out of 4 variables LRT *p* < 0.05; OP excluded = 0.087 ± 0.023, 3 out of 3 variables LRT *p* < 0.05) (*20*).

**Fig. 2.**
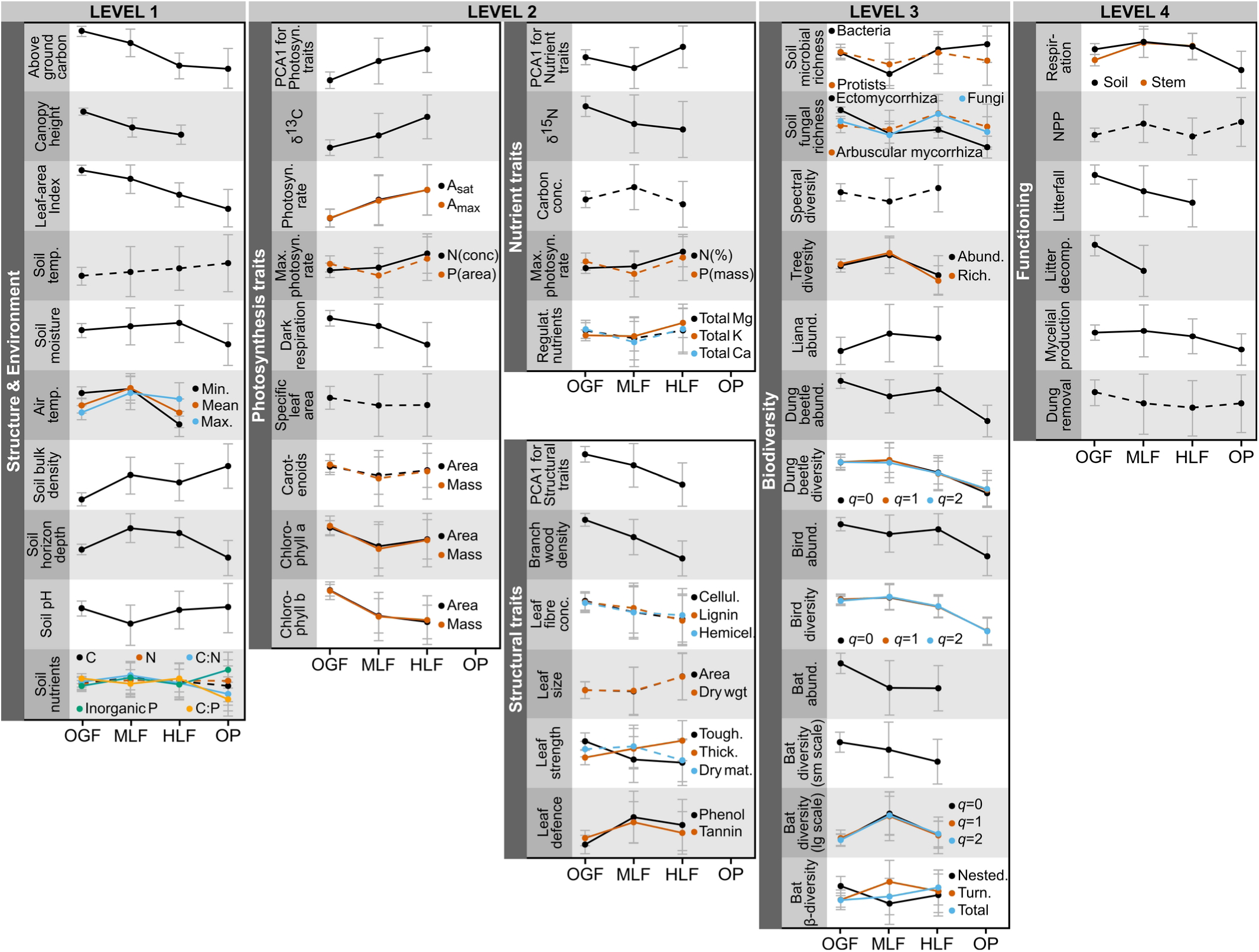
Changes in different categories of the measured response variables across the disturbance gradient when old-growth forest (OGF) is moderately logged (MLF), highly logged (HLF) and converted to oil palm plantation (OP). Points show z-score standardized means (± 95% C.I.). Sample sizes ranged from 27 to 373,968 depending on the dataset. Line type indicates whether a model with disturbance was significantly different from a null model with no disturbance (solid lines = significant, *p* < 0.05; dashed lines = non-significant). Tree traits (level 2) were analyzed individually (Fig. S3) as well as in combination via the first axis of a PCA (red backgrounds).

There was high variability in the observed patterns of responses to the disturbance gradient (Fig. 2). While some variables showed simple, monotonic responses (e.g., estimates of biomass carbon stocks decreased with disturbance, while frequency of photosynthetic traits associated with earlier successional species increased), other variables responded in a more complex manner (e.g., stem respiration increased in MLF but decreased to levels lower than in OGF in OP). Some patterns were also scale-dependent. For example, bat species richness decreased linearly across the disturbance intensity gradient at fine scales, but was highest in MLF at coarse scales, perhaps driven by an increase in community turnover and influxes of disturbance-adapted species in logged forest (*16*). Overall, different impacts of land-use change were heterogeneous, often non-linear and frequently not strongly correlated. Therefore, the impacts of logging and conversion defy simple interpretations, most likely due to a mixture of complex interactions, feedbacks and system redundancy.

## RESPONSES ALONG THE DISTURBANCE GRADIENT

To investigate the relative robustness of the different variables to disturbance intensity, we used sequential statistical contrasts to determine the stage along the disturbance gradient at which each variable was most affected: the initial logging of old-growth forest, further rounds of logging, or conversion to oil palm. The variables showed a wide range of responses, but with some ecological levels responding in broadly similar ways. Structural and environmental components of the forest (level 1) were generally more sensitive to a moderate degree of logging (Figs. 3, S2). This was especially the case for variables directly altered by the logging process itself, such as soil bulk density that was compacted by machinery (*21*), and above-ground carbon stock that was reduced by timber removal (*22–24*). This indicates that impacts on variables at level 1 are likely due to the direct effects of the timber removal and the conversion process, even several decades after they have taken place. Traits of the mature tree community (level 2) exhibit major changes consistent with the effects of selective logging (Fig. S3) (*18*, *25*), which actively targets tree species with the most commercially desirable characteristics. Removing individuals of these species effectively reduces the incidence of traits such as structural features that aid longevity (*25*), while increasing the incidence of traits such as high photosynthetic rates and rapid growth (*22*), which are associated with species of low commercial value and early successional species that colonize open areas following logging.

**Fig. 3.**
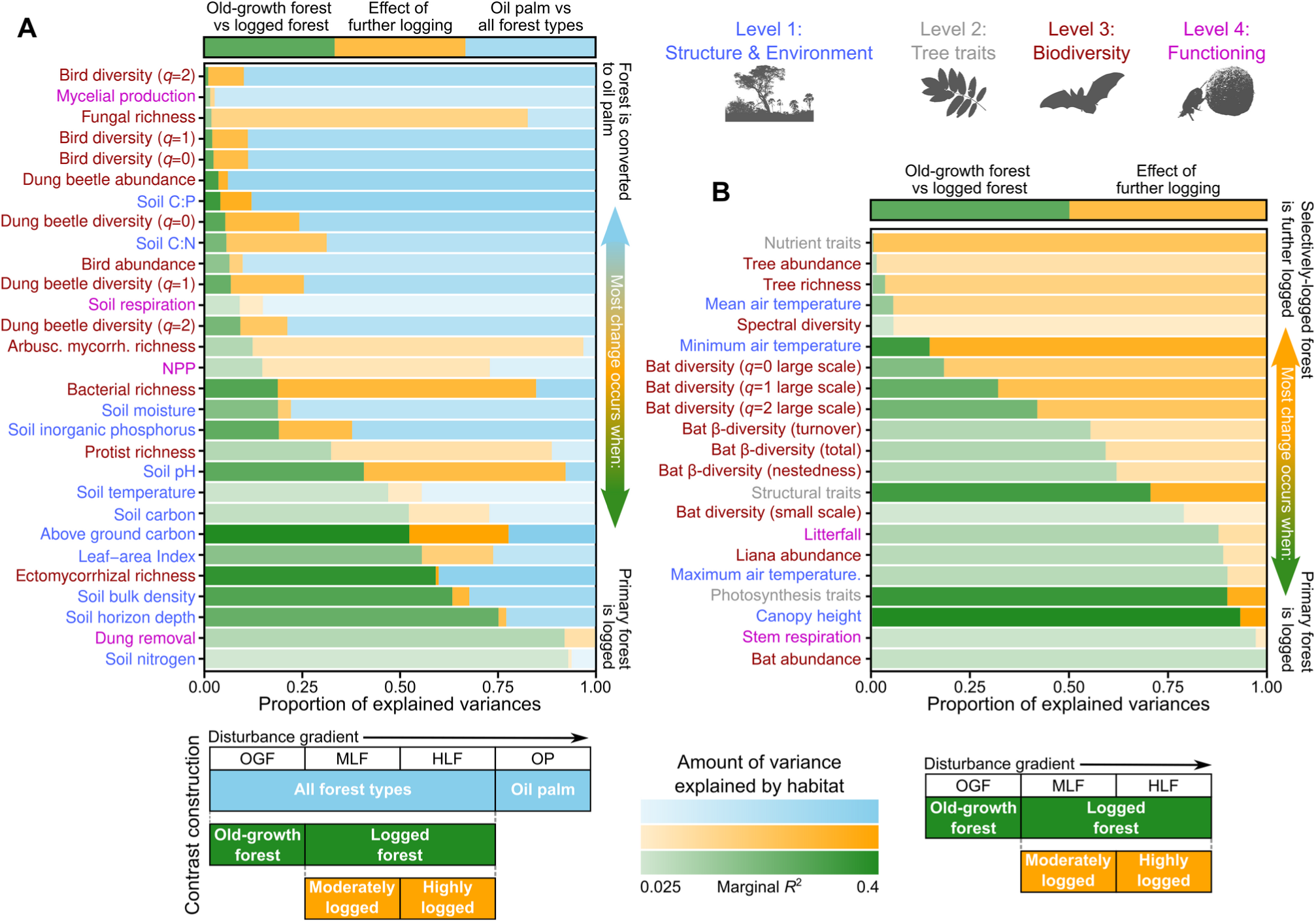
Impacts of different degrees and types of disturbance when old-growth forest (OGF) is moderately logged (MLF), highly logged (HLF) and converted to oil palm plantation (OP). The overall effect of disturbance was partitioned into a) three (for datasets that did sample in oil palm), or b) two (for datasets that did not sample in oil palm) or single degree of freedom contrasts that compare: 1) the effect of logging old-growth forest (old-growth *vs* logged forest: OGF *vs* MLF-HLF; green bars); 2) further logging of moderately logged forest (MLF *vs* HLF; orange bars); and 3) converting forest to oil palm plantation (oil palm *vs* the combined forest types: OP *vs* OG-MLF-HLF; blue bars). Sample sizes ranged from 27 to 373,968 depending on the dataset. The transparency for each variable is inversely related to its explanatory power (the marginal *R*^2^ from the linear mixed-effects model). Response variables are categorized into ecological levels: those related to the forest structure and environment (level 1; blue text); those related to biodiversity (level 3; dark red text); and those related to ecosystem functioning (level 4; dark green text), and are ordered by the size of the effect of logging OGF (variables at the bottom are mostly affected when OGF is logged, whereas those at the top are mostly affected when forest is converted to oil palm).

In contrast, biodiversity components (level 3) were mostly altered by the conversion of logged forest to oil palm (Figs. 3, S4) (*6*, *10*, *26*). This was particularly true of taxonomic groups at higher trophic levels, such as birds (*10*, *23*), where increased mobility and behavioral plasticity may buffer their sensitivity to change until conditions become drastically altered (*27*). In the highly disturbed conditions of oil palm plantations, the major changes in plant food resources, reduced structural complexity, and shift to hotter, drier, and more variable microclimatic conditions, likely resulted in the reduced abundance and diversity of many taxa, and a community composed of disturbance-tolerant species (*28*). The diversity of ectothermic groups, such as dung beetles, which can be particularly responsive to changes in microclimate (*29*), showed a slightly increased sensitivity to the initial impacts of logging relative to endothermic taxa, such as birds and bats. The richness of soil microorganisms showed the greatest sensitivity to logging, although there were both negative (ectomycorrhizal fungi (*30*, *31*)) and positive (bacteria (*32*)) responses to disturbance. The impact on ectomycorrhizal fungi is likely to be particularly important for conservation and restoration perspectives, given their role in supporting canopy-dominant dipterocarps (*33*).

Finally, ecosystem functions (level 4; Figs. 3, S4) showed the weakest and most variable patterns (marginal *R*^2^ values shown by high transparency of bars in Fig. 3, Table S6). For example, rates of dung removal were maintained in oil palm plantations, even though dung beetle richness and abundance decreased significantly with disturbance (*34*), with a small number of disturbance-tolerant species increasing their contribution to dung removal in disturbed habitats (*35*). Such functional redundancy and compensation may confer greater resilience (here, the ability to both resist and recover from change) of ecosystem functions to disturbance (“insurance hypothesis” (*36*) or “portfolio effect” (*37*)). For example, previous research at these study sites found that, while certain taxonomic groups that dominated litter decomposition and seed and invertebrate predation in old-growth forests declined along the logging gradient, different taxonomic groups compensated by increasing their contribution, and so maintained those ecosystem functions at similar levels (*27*). The relative stability of ecosystem functions may also be related to the large spatial extent and connectivity of the habitat blocks investigated, which can enhance the relationship between biodiversity and ecosystem functions and services (*38*).

## IMPLICATIONS FOR TROPICAL FOREST CONSERVATION

Our findings increase understanding of the ecosystem-wide impacts of habitat change in the tropics, with implications for land-use management and restoration. Although our large-scale study shares the standard limitations of the space-for-time approach to observational data, by adopting a unified framework that considers all measured variables, we reveal that selective logging and forest conversion to oil palm plantation have different environmental impacts, and that these vary depending on which aspects of the ecosystem are considered. Even a single type of land use can have a range of impacts when the environment is assessed comprehensively to encompass its abiotic and biotic structure, biological diversity, and the multiple ecosystem functions and services that it provides. This emphasizes the importance of considering a broad range of ecological properties when making land-management, conservation, and research decisions.

Our finding that factors associated with forest structure and environment (level 1) are highly sensitive to disturbance shows that even low intensity logging will result in changes in these characteristics, and highlights the importance of maintaining areas of intact, undisturbed forest. The large negative effects on biodiversity when forest is converted to oil palm are consistent with findings from other studies in the region (*23*, *26*), and confirm the value of disturbed forest for the maintenance of high overall biodiversity at the landscape scale (*39*). Therefore, while preserving areas of remaining old growth forests is important for conserving unique aspects of their biodiversity and functioning, protecting logged forest can also contribute to maintaining biodiversity and ecosystem functioning relative to landscapes with higher levels of conversion to agriculture. This validates an increasing focus within tropical agricultural systems of maintaining forest in sensitive areas within plantations, such as steep slopes and river margins (e.g., as highlighted by the Roundtable on Sustainable Palm Oil (RSPO) (*40*)), where it can support both biodiversity (*41*) and ecosystem processes (*42*). The reduction in some taxa, such as birds and ectomycorrhizal fungi, and some ecosystem functions, such as mycelial production, within oil palm also has implications for crop management, and could affect nutrient cycling and predator control (*20*).

Understanding at which points on the deforestation gradient biodiversity and associated ecosystem functions are most affected is important for identifying priority habitats for conservation and restoration (*39*, *43*, *44*), and can aid decision-making in these complex, multi-use landscapes (*45*, *46*). However, it is important to consider multiple facets of these tropical environments in order to avoid the risk of unintended consequences possible from more narrow assessments. This study provides an initial comprehensive synthesis and overview of the responses of a tropical forest ecosystem to degradation and deforestation. However, despite the breadth of ecosystem properties investigated, this study represents only a single area and ecosystem type: lowland tropical forest. Future research should establish whether these responses are consistent across other tropical landscapes and in relation to the wider range of land-use changes seen across the global tropics.

## Supporting information

Materials, methods and further results are available as supplementary materials

## Acknowledgments

We thank Sabah Forestry Department, Sabah Foundation and Universiti Malaysia Sabah. We thank Dan Lunn of Oxford University Statistical Consultancy for advice.

We are grateful to the South East Asia Rainforest Research Partnership (SEARRP) for logistical support, and Dr Chey Vun Khen for his support and advice in the initial SAFE Project set-up.

Fieldwork was supported by Rasizul Bin Sahamin, Joshua Blackman, Genevieve Durocher, Ryan Gray, Walter Huaraca Huasco, Unding Jami, Rostin Jantan, Alexander Karolus, Raina Manber, Toby Marthews and SEARRP staff, including Amir, Anis, Austin, David, Didy, Dino, Lizzie, Johnny, Kiki, Loly, Mudin, Noy and Zul. We thank Yayasan Sabah, Maliau Basin Management Committee, Danum Valley Management Committee, the State Secretary, Sabah Chief Minister’s Departments, the Malaysian Economic Planning Unit and the Sabah Biodiversity Council for permission to conduct research.

## Funding

Data were collected and analyzed as part of the BALI (Biodiversity And Land-use Impacts on tropical ecosystem function) and LOMBOK (Land-use Options for Maintaining BiOdiversity & eKosystem functions) projects within the Human-modified Tropical Forests Programme funded by NERC (NE/K016377/1, NE/K016261/1, NE/K016148/1, NE/K016407/1).

Collection of dung beetle data was also supported by a British Ecological Society Small Ecological Project Grant, No.: 3256/4035, and the Varley-Gradwell Travelling Fellowship in Insect Ecology to EMS.

Collection of bird data was supported by NERC (NE/I028068/1) to JAT.

Collection of bat data was supported by a Bat Conservation International Student Research Scholarship to DHB.

TR acknowledges support from the European Research Council under the European Union’s Horizon 2020 research and innovation programme (grant agreement No 865403).

Carbon cycle work was also supported by European Research Council Advanced Investigator Grant, GEM-TRAIT (321131) to YM.

The work of the Leverhulme Centre for Nature Recovery is made possible thanks to the generous support of the Leverhulme Trust.

The SAFE Project is funded by the Sime Darby Foundation and RME by the NOMIS Foundation.

## Author contributions

Paper conceived, and initial draft prepared by CJM, ECT, AH. CJM performed all analyses. All authors contributed to further drafts and provided final approval. Data collection and processing was carried out by BWB, BB, SB, RSC, DMOE, DPE, DH-B, PJ, VK, UKH, SM, DTM, SLM, MMP, MHN, TR, SJBR, EMS. Principal investigators of individual projects were HB, DFRPB, AYCC, ELC, DAC, ZGD, DPE, DJ, PK, YM, NM, RN, NJO, SJR, MJS, JAT and MW. The BALI and LOMBOK projects were planned by the project teams led by PIs YAT and OTL, and the SAFE Project established and led by GR and RME.

## Data and materials availability

Markdown documents containing R code and their outputs for the exploration of data, analysis and presentation of results for all 82 variables used in the study, along with the processed z-score standardised data, are available at https://zenodo.org/records/13161799. The DOIs for archived versions of the raw datafor all datasets are listed in the methods and Tables S2-5.

## Materials and Methods

### Location and study design

All datasets come from lowland tropical forest or oil palm plantation in Sabah, Malaysian Borneo (Fig. 1). Sampling within old-growth forest occurred in Maliau Basin and Danum Valley Conservation Areas. These were compared with nearby areas of logged forest, including those that are part of the Stability of Altered Forest Ecosystems (SAFE) Project (*14*) — a long-term, large-scale study of forest degradation and fragmentation. The SAFE Project area has undergone multiple rounds of selective and clearance logging in preparation for oil palm planting. Sampling in oil palm occurred within mature plantations adjacent to the SAFE Project.

Due to the large breadth of studies examined, datasets utilized multiple sampling sites and structures, including points, transects and remotely-sensed data (Table S1). Detailed descriptions of each dataset’s sampling design and data characteristics, as well as all analytical steps and outputs, can be found within the RMarkdown document for each analysis at https://zenodo.org/records/13161799 (*47*), and summarised in Tables S2-6. The majority of datasets (77 %) were collected within a network of 1 ha forest plots used for investigating carbon dynamics (*22*), hereafter ‘carbon plots’. We assigned carbon plots to three categories following Riutta *et al.* 2018 (*22*) (see Table 1 in that publication) along a disturbance gradient: old-growth forest (OGF; n = 4), moderately logged forest (MLF; n = 2) and highly logged forest (HLF; n = 3), as well as a smaller plot of 60 × 60 m in oil palm (OP; *n* = 1). Individual studies used different combinations of the available carbon plots, outlined under the column ‘No. largest groups’ in Table S1. Studies using other sampling schemes were placed into the appropriate categories to enable comparison, as outlined in each RMarkdown document.

OGF plots had a larger basal area of mature trees (basal area of trees > 10 cm (m^2^ ha): OGF = 30.6 – 41.6; MLF = 19.3 – 19.6; HLF = 6.81 – 13.9), a more closed canopy (mean canopy gap fraction (%): OGF = 7.04 – 11.3; MLF = 11.2 – 12.8; HLF = 12.2 – 15.0), more large trees (number of large trees (number of trees per ha with DBH > 50 cm): OGF = 26 – 56; MLF = 10 – 11; HLF = 0 – 6) and the lowest proportion of pioneer trees (% basal area composed of pioneer trees: OGF = 0.1 – 1.7; MLF = 6.9 – 21.5; HLF = 28.1 – 57.2) compared to other habitats (all values taken from Table 1 in (*22*)).

The SAFE Project site was logged in 1978, removing around 113 m^3^ ha^−1^. Areas of heavily logged forest were logged again in the late 1990s to the early 2000s in three rounds (salvage logging with a view to conversion), resulting in a cumulative extraction rate of 66 m^3^ ha^−1^ (*16*). Areas of moderately logged forest were logged twice at approximately the same times, but with around 37 m^3^ ha^−1^ removed during the second rotation (*16*). Finally, the area was further salvage-logged between 2013 and 2016, with the logging front gradually moving across from the south to the north of the SAFE area, removing any remaining trees, but with small forest patches on slopes being maintained (*48*). Oil palm plantations were established in 2000 and 2006.

For analyses, each study was decomposed into appropriate spatial hierarchies depending on its sampling design. For example, studies within the 1 ha carbon plots were divided into 25 subplots of 20 × 20 m (400 m^2^), and these subplots were spatially clustered into groups of 5 quadrats of 2000 m^2^ (Fig. S1). Other studies used the SAFE Project sampling design, where sites are situated at the apices of a fractal series of triangles (*14*). Each triangle, of area 9,000 m^2^, consists of 27 sampling points. Three triangles are grouped together into blocks (150,000 m^2^). Where a study used neither carbon plots nor SAFE Project fractals, points were grouped spatially as appropriate. Details for each dataset, including the number and grain size of spatial groups, are outlined in Table S1, and more detailed descriptions of the processing carried out on each dataset can be found in the accompanying RMarkdown document.

### Datasets

A total of 82 datasets were compiled (*49–62*). In many cases, individual studies were analysed at multiple facets and levels. For example, a study sampling bats was analysed as eight separate datasets examining multiple facets of diversity at multiple scales. Datasets were categorized into broad hierarchies representing ecological levels, where each higher strata can be considered an emergent feature of the lower levels. Tables S3-6 give brief descriptions of data collection and preparation for each dataset with RMarkdown documents outlining the process in detail.

STRUCTURE & ENVIRONMENT (level 1) — the abiotic environment and forest structure, such as soil properties, microclimate and above-ground carbon. These are variables that are directly affected by logging through direct manipulation of the forest structure with tree removal, or soil compaction by machinery used to extract timber (*21*).

TREE TRAITS (level 2) — traits related to life-history strategies of trees present within plots. Changes to the distribution of these traits is a direct consequence of the species selected for removal, as well as the subsequent growth and recruitment of early-successional species in open areas generated after mature trees were extracted. Traits have been categorized into three groups (*25*): a) structural traits related to investment in stability and defence; b) photosynthesis traits related to leaf photosynthetic potential and leaf longevity; and c) nutrient traits related to concentrations of key nutrients. Oil palm monocultures were not investigated for tree traits. Descriptions of the individual traits can be found in Table S3. Traits within each of the three groups were presented individually and also summarised via the first axis of a PCA ( Fig. 2; red backgrounds in and Fig. S3). All tree traits were collected as part of a single campaign (*25*) using the same locations and tree individuals over a single sampling period.

BIODIVERSITY (level 3) — below-and above-ground biodiversity at multiple trophic levels (bacteria, protists and fungi, trees, beetles, bats and birds), as well as functional diversity (spectral diversity). Species are dependent upon forest structure, microclimate and the tree species remaining or establishing post-logging. Where possible we explored multiple facets of diversity (α-diversity, β-diversity and abundance) at multiple scales (Table S1). For example, bat data were analysed as abundance and species richness at the finest sampling grain sizes, but we also aggregated data to coarser spatial grain sizes. Where possible, we analysed α-diversity as three components using effort-standardised Hill numbers (*19*), as species richness (*q* = 0), Shannon diversity (*q* = 1) and Simpson’s diversity (*q* = 2), which explores changes to the structure of each community by either excluding species relative abundances (when *q* = 0) through to considering species in proportion to their relative abundance (when *q* = 1) and finally estimating the effective number of dominant species in the community at a given site (when *q* = 2). For tree richness, where we could identify all individuals (≥ 10 cm DBH) within each subplot, we did not standardise by effort. We also decomposed total presence-based Sorensen’s β-diversity into turnover and nestedness components (*63*, *64*), in order to examine whether differences between communities over space are due to species replacement from one site to another, or community subsetting due to richness differences respectively.

FUNCTIONING (level 4) — ecosystem functions that are driven by the abundance and composition of the relevant communities. Functions can be relatively resilient (which here incorporates both the degree of resistance to disturbance, as well as its rate of recovery) to forest degradation as the loss of a dominant functional group can be replaced by another (*27*). We focussed on lower-level functions related to productivity (soil and stem respiration; net primary productivity) and nutrient cycling (leaf litter fall and decomposition; dung removal).

### Statistical analyses

For reproducibility and transparency, all data exploration, manipulation and analytical procedures were carried out with RMarkdown scripts, which are made available at https://zenodo.org/records/13161799 (*47*). All data manipulation, analyses and visualisation were carried out in R 4.0+ (*65*) using the packages betapart (*66*), cowplot (*67*), dismo (*68*), dplyr (*69*), extrafont (*70*), ggplot2 (*71*), gridExtra (*72*), gtable (*73*), iNEXT (*19*, *74*), lattice (*75*), lme4 (*76*), magrittr (*77*), MuMIn (*78*), optimx (*79*, *80*), readr (*81*), rgeos (*82*), rmarkdown (*83*), sp (*84*, *85*), tidyr (*86*), terra (*87*), vegan (*88*). Specific versions of packages used for each dataset are recorded at the end of each RMarkdown document. To allow for comparability between many datasets with multiple data types, analyses were kept relatively simple, using only disturbance category as a predictor variable and without other explanatory co-predictors. We therefore only examine patterns along the disturbance gradient and do not explore the processes behind them.

For detailed investigations examining the causes of these patterns, please see the individual publications for each dataset outlined in Tables S3-5. Although we follow convention in presenting estimates of statistical significance, we focus on effect sizes, confidence intervals and explained variance (*89*).

### Change along the disturbance gradient

We examined the change in values of each dataset along the disturbance gradient (included as a categorical variable with up to four levels depending on the dataset: OGF, MLF, HLF and OP), using linear mixed-effects models to account for the different hierarchical sampling schemes of datasets, using the lme4 package, v1.1_27-31 (*76*). Hierarchical components were included as nested random effects as appropriate for the study, as well as a separate date component if required for time-series data with temporal replicates (Table S1: column ‘Temporal replicates’). Only disturbance, as a non-ordered categorical explanatory variable, was included as a fixed effect (with 3 or 4 levels: OGF, MLF, HLF, OP). The generic model (using the R language implementation of the Wilkinson and Rogers syntax (*90*) was:

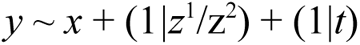

where: *y* is one of the 82 forest property response variables listed in Tables S2-5; *x* is a fixed factor for disturbance; *z*^1^ and *z*^2^ are random factors denoting nested spatial sampling structures as appropriate to the dataset, and *t* is a variable for time (e.g date) if a data set has repeated samples through time. Exact model formulations were defined by the sampling structures of each dataset, and can be seen in the RMarkdown document outlining the analysis in each case.

Linear mixed-effects models are intrinsically more conservative than other statistical models due to the shrinkage of coefficients, and therefore reduce the chances of false positives (*91*). Rather than examining individual datasets for significance, we focus on the overall number of significant relationships identified in each ecological level in relation to the number of false positives we would expect from chance, given a *p*-value of 0.05.

We first inspected the data and applied the most appropriate transformation if required. If a transformation was explored, we also constructed a model with non-transformed data for comparison. Both sets of data, transformed and non-transformed, were z-score standardised for analysis. We controlled for differences in the variance between disturbance categories through the addition of weights as the inverse of the variance, calculated separately for transformed and non-transformed data if necessary.

We assessed model assumptions using standard graphical assessments including plots of the fitted values against residuals (the Tukey-Anscombe plot) and square-root transformed residuals (the scale location plot), plots of the residual variance within disturbance categories and quantile-quantile plots. We selected the model using transformed data if it was a visual improvement over the model with non-transformed data, otherwise the model using non-transformed data was retained. We calculated 95% confidence intervals of the best model using parametric bootstrapping with 500 permutations, or by using the Wald statistic, where bootstrapped models failed to converge.

We determined the significance of the model for each dataset using ANOVA by comparison to a null model incorporating the same random effects structure, but without disturbance category as a fixed effect, refitting the models with log-likelihood (ML) rather than restricted log-likelihood (REML). We calculated the percentage of variance explained by disturbance through marginal (*R*^2^_cond._) and conditional (*R*^2^_marg._) coefficients of determination using the MuMIn package (*78*). The *p*-values, marginal *R*^2^ and effects sizes of the selected models of all variables are presented in Table S6.

As some datasets in levels 1 and 3 were analysed across multiple facets (for example, bats were examined for α-diversity and β-diversity at multiple spatial scales), we also performed a sensitivity analysis controlling for dataset identity. Response variables were grouped according to Table S1 and then we randomly drew one variable from each group and calculated the number of significant and non-significant variables. We repeated this 1000 times and calculated the mean proportion of significant to non-significant variables. The proportions when controlling for dataset were very similar compared to when using all response variables (level 1: 81.25 % significant for all variables, 82.48 % significant when randomising; level 3: 88 % significant for all variables, 81.75 % significant when randomising).

### Contrasts between disturbance categories

We examined where the changes in our observed values occurred along the disturbance gradient using model contrasts, which decomposes the explained variance into pairwise comparisons between the four habitat types. The proportions of explained variance were visually compared using *R*^2^ as an indication of the total variance explained by disturbance alone (i.e. there may be cases where a contrast may explain a high proportion of the total explained variance in the best model, but as a whole explain only a small amount of the overall variance between disturbance categories). The exact calculation of the contrasts depended upon whether the dataset had samples within OP or the three forest habitats only. One dataset (litter decomposition) that had only sampled OGF and MLF was excluded from this part of the analysis.

For datasets that had sampled OP, we examined two contrasts sequentially. The first examined the difference between OP and all forest habitats (OGF, MLF and HLF) by generating a new categorical variable (‘contrastOP’). The second contrast examined the difference between OGF against the disturbed forest habitats (MLF and HLF) using a second categorical variable (‘contrastOGF’). The remainder is the effect of re-logging of selectively-logged forest (MLF vs HLF). Contrasts were fitted with the two new categorical variables in order, followed by the remaining disturbance variable, retaining the random effect structure (i.e. a fixed effects structure of ∼ contrastOP + contrastOGF + Disturbance). We then calculated the variance explained by each contrast, relative to the total explained variance as the sums of squares for that contrast as a proportion of the total sums of squares explained by all contrasts of the Analysis of Variance table. This produced three variance components: 1) the effect of converting all forest types to oil palm (OP-OGF, OP-MLF and OP-HLF); 2) the effect of logging old-growth forest to medium-or highly-logged forest (OGF-MLF and OGF-HLF); and 3) the effect of converting medium-to highly-logged forest (MLF-HLF).

For datasets that had not sampled OP, we eliminated the first contrast comparing OP to all forest habitats (i.e. a fixed effects structure of ∼ contrastOGF + Disturbance). These datasets therefore only compared 1) the effect of logging old-growth forest to medium-or highly-logged forest (OGF-MLF and OGF-HLF); and 2) the effect of converting medium-to highly-logged forest (MLF-HLF).

