## Supplementary material for "Logging alters tropical forest structure, while conversion to agriculture reduces biodiversity and functioning": Materials, methods and further results are available as supplementary materials

**This PDF file includes:**

Materials and Methods
Figs. S1 to S4
Tables S1 to S6

**Other Supplementary Materials for this manuscript include the following:**

Markdown outputs outlining the analyses of all 82 variables used in the study, along with the processed z-score standardised data, are available at https://zenodo.org/records/13161799. DOIs for archived versions of the raw data for all datasets are listed in the methods and in Table S2-5.

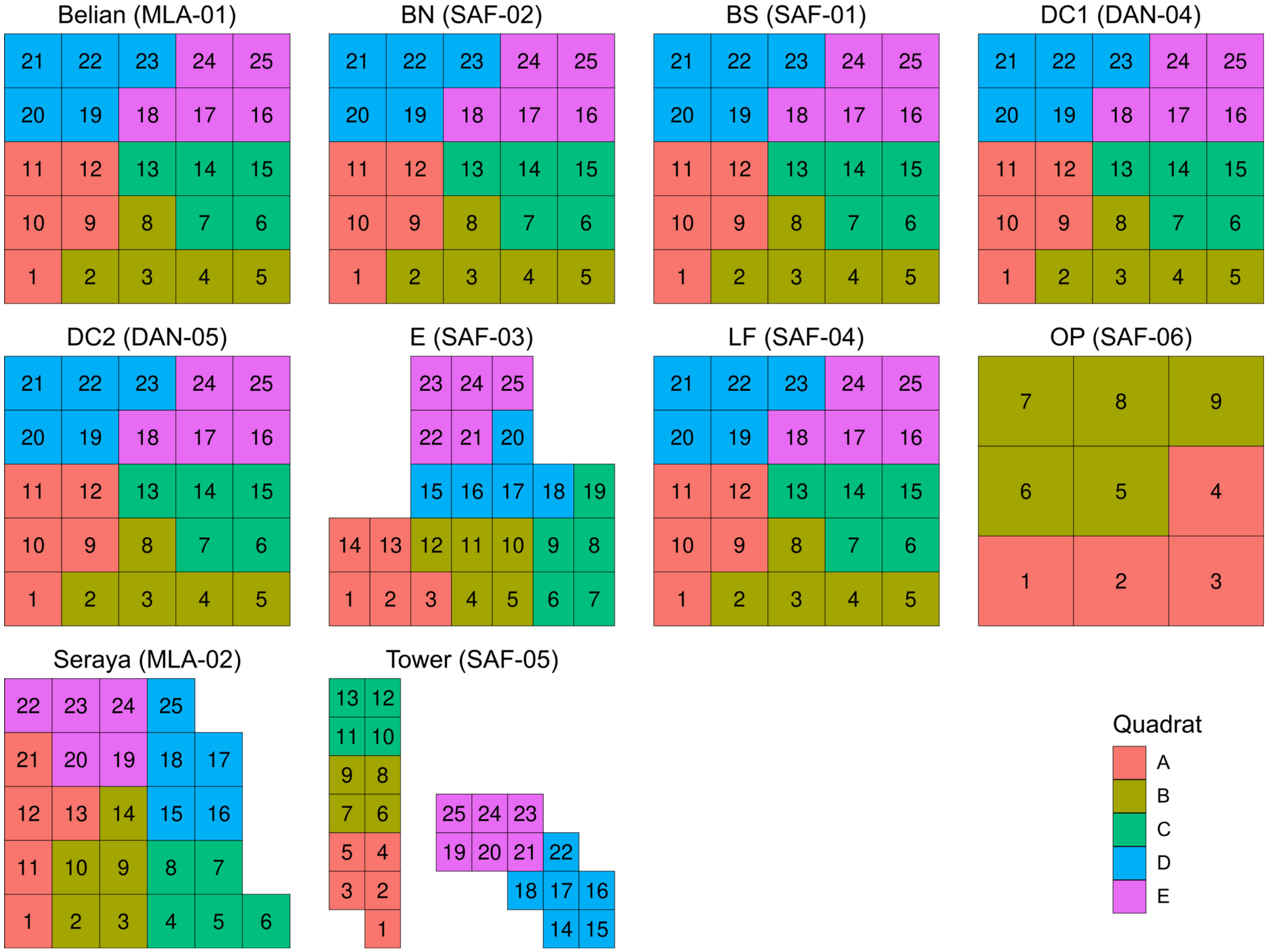

**Figure S1 – Schematics for the spatial clustering used for the 1 ha (10,000m$^2$) carbon plots** (3,600 m$^2$ for the oil palm plot). Plots were split into 20 × 20 m (400 m$^2$) subplots. Subplots were clustered into groups of 4-6 quadrats (1,600 – 2,400 m$^2$). Note that each carbon plot is represented in schematic form and are not drawn at the same scale. In reality, all subplots across all carbon plots are of equal area.

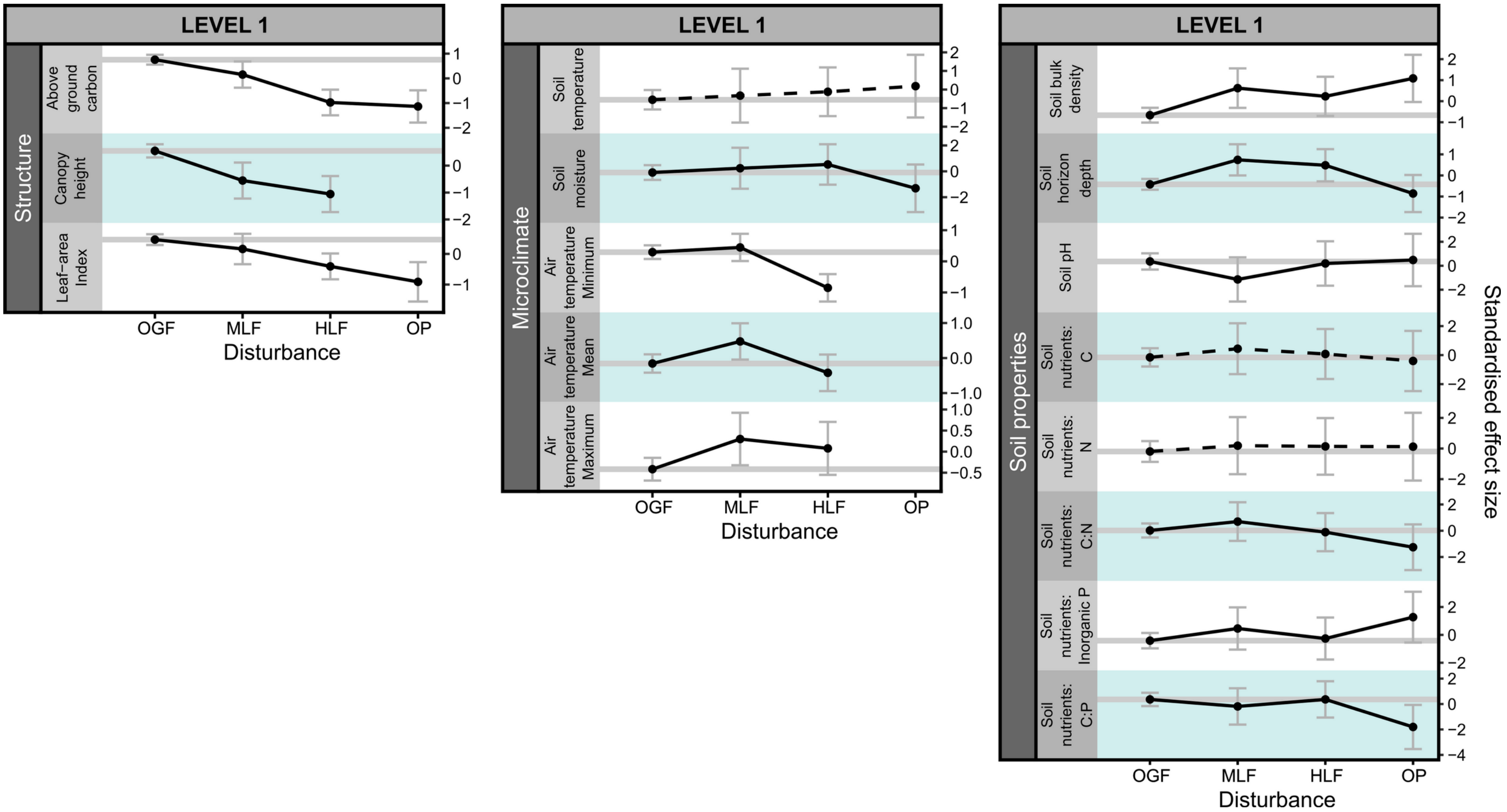

**Figure S2 – Changes in different categories of the measured response variables in ecological level 1 (forest structure, microclimate and soil properties) across the disturbance gradient when old-growth forest (OGF) is moderately logged (MLF), highly logged (HLF) and finally converted to oil palm plantation (OP).** Points show z-score standardized means (± 95% C.I.). Line type indicates whether a model with disturbance was significantly different from a null model with no disturbance (solid lines = significant; dashed lines = non-significant; significance threshold = $p < 0.05$). Grey lines indicate mean levels in OGF.

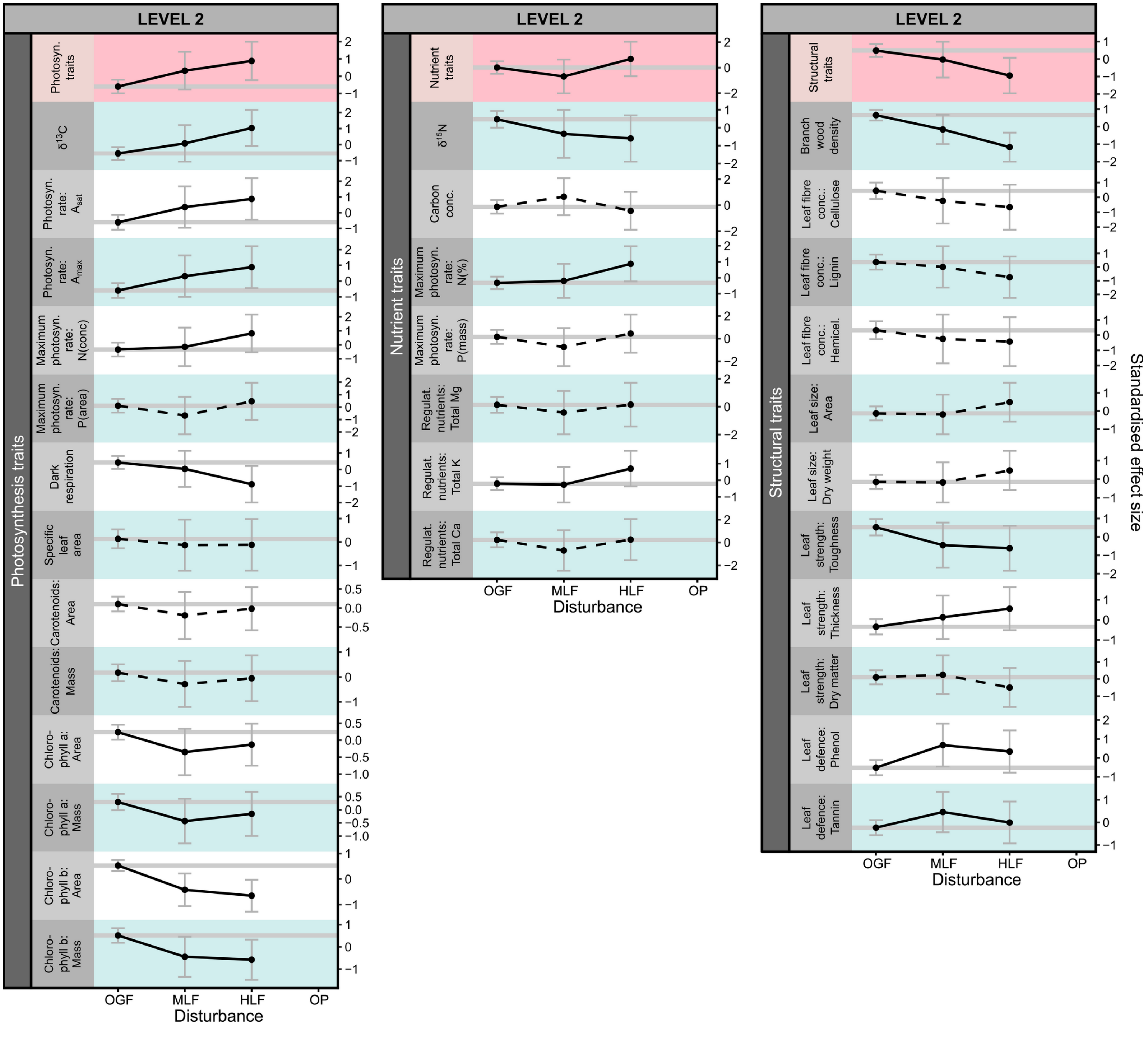

**Figure S3 – Changes in different categories of the measured response variables in ecological level 2 (tree traits categorised into structural, nutrient and photosynthesis traits) across the disturbance gradient when old-growth forest (OGF) is moderately logged (MLF), highly logged (HLF) and finally converted to oil palm plantation (OP).** Points show z-score standardized means (± 95% C.I.). Line type indicates whether a model with disturbance was significantly different from a null model with no disturbance (solid lines = significant; dashed lines = non-significant; significance threshold = $p < 0.05$). Grey lines indicate mean levels in OGF. Traits were analysed individually as well as in combination via the first axis of a PCA (red backgrounds).

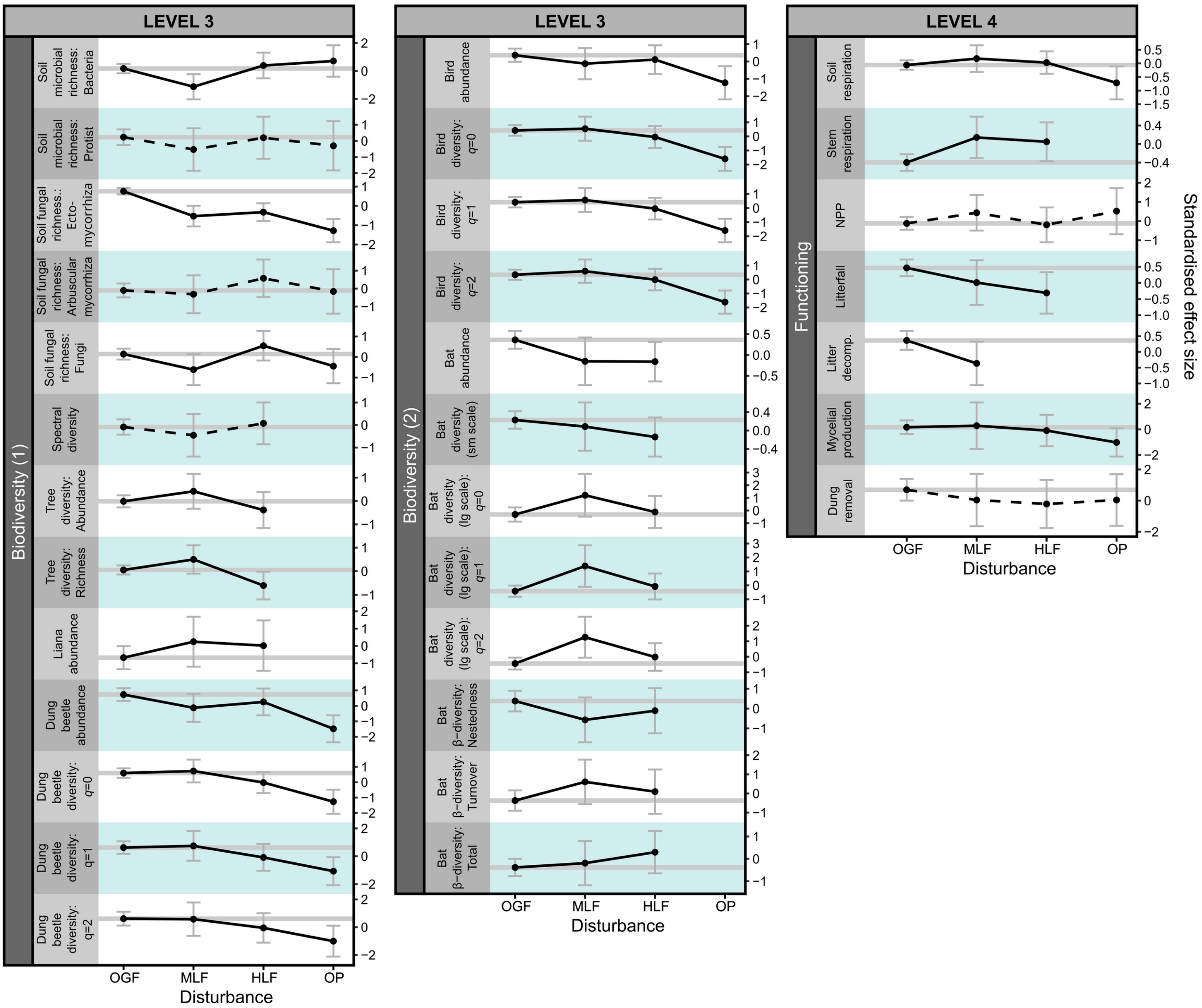

**Figure S4 – Changes in different categories of the measured response variables in ecological levels 3 (biodiversity) and 4 (functioning) across the disturbance gradient when old-growth forest (OGF) is moderately logged (MLF), highly logged (HLF) and finally converted to oil palm plantation (OP).** Points show z-score standardized means (± 95% C.I.). Line type indicates whether a model with disturbance was significantly different from a null model with no disturbance (solid lines = significant; dashed lines = non-significant; significance threshold = $p < 0.05$). Grey lines indicate mean levels in OGF.

**Table S1 – Summary of characteristics of each dataset**. Outlined are whether the study sampled in old-growth forest (OGF), moderately logged forest (MLF), highly logged forest (HLF) or oil palm plantation (OP), the total number of sampling points (*n*), year(s) that data were collected, and the total area and number of the smallest and largest spatial groupings of the spatial hierarchy of the sampling design for each study, which were specified as random effects in linear mixed-effects models. Some datasets also used time-series data, in which case the temporal replicates were also specified as a date in the random effects. Datasets that used the same sampling events have been combined into single rows, and are described in more detail in Tables S3-6 and in the RMarkdown documents outlining the processing of each dataset in detail.

| **Dataset** | **OGF** | **MLF** | **HLF** | **OP** | ***n*** | **Largest spatial grouping** | **No. largest groups** | **Area of largest groups** | **Smallest spatial grouping** | **No. smallest groups per largest group** | **Area of smallest groups** | **Year(s)** | **Temporal replicates** |
|---|---|---|---|---|---|---|---|---|---|---|---|---|---|
| ***Structure & Environment*** | | | | | | | | | | | | | |
| Above ground carbon | X | X | X | X | 234 | Carbon plots | 10 | 10,000 $m^2$ (3,600 $m^2$ for OP) | Subplot | 25 (9 for OP) | 400 $m^2$ | 2011-2016 | No |
| Canopy height | X | X | X | | 373,968 | Carbon plots | 8 | 10,000 $m^2$ | Subplot | 25 | 400 $m^2$ | 2014 | No |
| Leaf-area Index | X | X | X | X | 2,429 | Carbon plots | 10 | 10,000 $m^2$ (3,600 $m^2$ for OP) | Subplot | 25 (9 for OP) | 400 $m^2$ | 2013-2016 | Yes |
| Soil temperature | X | X | X | X | 3,329 | Carbon plots | 10 | 10,000 $m^2$ (3,600 $m^2$ for OP) | Subplot | 25 (16 for OP) | 400 $m^2$ | 2015-2017 | Yes |
| Soil moisture | X | X | X | X | 222 | Carbon plots | 9 | 10,000 $m^2$ (3,600 $m^2$ for OP) | Subplot | 5 | 400 $m^2$ | 2015 | No |
| Air temperature (Minimum, Mean, Maximum) | X | X | X | | 4727 | Carbon plots | 3 | 10,000 $m^2$ | Subplot | 24-25 | 400 $m^2$ | 2015 | Yes |
| Soil properties (bulk density, horizon depth, pH) | X | X | X | X | 222-225 | Carbon plots | 9 | 10,000 $m^2$ (3,600 $m^2$ for OP) | Subplot | 5 | 400 $m^2$ | 2015 | No |
| Soil nutrients (C, N, C:N, Inorganic P, C:P) | X | X | X | X | 225 | Carbon plots | 9 | 10,000 $m^2$ (3,600 $m^2$ for OP) | Subplot | 5 | 400 $m^2$ | 2015 | No |
| ***Tree traits*** | | | | | | | | | | | | | |
| Photosynthesis traits | X | X | X | | 198 | Carbon plots | 8 | 10,000 $m^2$ | Subplot | 24-25 | 400 $m^2$ | 2015 | No |
| Nutrient traits | X | X | X | | 198 | Carbon plots | 8 | 10,000 $m^2$ | Subplot | 24-25 | 400 $m^2$ | 2015 | No |
| Structural traits | X | X | X | | 198 | Carbon plots | 8 | 10,000 $m^2$ | Subplot | 24-25 | 400 $m^2$ | 2015 | No |

| | | | | | | | | | | | | | |
|---|---|---|---|---|---|---|---|---|---|---|---|---|---|
| ***Biodiversity*** | | | | | | | | | | | | | |
| Soil microbial richness (Bacteria, Protist) | X | X | X | X | 223-224 | Carbon plots | 9 | 10,000 $m^2$ (3,600 $m^2$ for OP) | Subplot | 5 | 400 $m^2$ | 2015 | No |
| Soil fungal richness (Ectomycorrhiza, Arbuscular mycorrhiza, Fungi) | X | X | X | X | 222-224 | Carbon plots | 9 | 10,000 $m^2$ (3,600 $m^2$ for OP) | Subplot | 5 | 400 $m^2$ | 2015 | No |
| Spectral diversity | X | X | X | | 673 | Carbon plots | 8 | 10,000 $m^2$ | Subplot | 3-23 | 400 $m^2$ | 2015 | No |
| Tree diversity (abundance and richness) | X | X | X | | 200 | Carbon plots | 8 | 10,000 $m^2$ | Subplot | 25 | 400 $m^2$ | 2016 | No |
| Liana abundance | X | X | X | | 390 | Block | 5 | ~23.5 ha | 2nd-order fractal points & Carbon plots | 1-13 | 10,000 $m^2$ | 2016 | No |
| Dung beetle diversity (abundance, q=0, q=1, q=2) | X | X | X | X | 176 | Block (9 2nd-order fractal points) | 15 | ~23.5 ha | Trio of 2nd-order fractal points | 3-13 | ~1.37 ha | 2011-2015 | No |
| Bird diversity (abundance, $q$=0, $q$=1, $q$=2) | X | X | X | X | 193 | Block (9 2nd-order fractal points) | 16 | ~23.5 ha | Trio of 2nd-order fractal points | 2-4 | ~1.37 ha | 2014-2017 | No |
| Bat small scale diversity (abundance, richness) | X | X | X | | 432-435 | Site | 8 | 37-86 ha | Group | 1-7 | 6-16 ha | 2015-2017 | No |
| Bat large scale diversity ($q$=0, $q$=1, $q$=2) | X | X | X | | 68 | Site | 8 | 37-86 ha | Cluster | 2-4 | 18-28 ha | 2015-2017 | No |
| Bat β-diversity (Nestedness, Turnover, Total) | X | X | X | | 55 | Site | 8 | 37-86 ha | Cluster | 1-4 | 18-28 ha | 2015-2017 | No |
| ***Functioning*** | | | | | | | | | | | | | |
| Respiration: Soil | X | X | X | X | 2,748 | Carbon plots | 10 | 10,000 $m^2$ (3,600 $m^2$ for OP) | Subplot | 25 (16 for OP) | 400 $m^2$ | 2015-2017 | Yes |
| Respiration: Stem | X | X | X | | 5,869 | Carbon plots | 9 | 10,000 $m^2$ | Subplot | 19-25 | 400 $m^2$ | 2011-2016 | Yes |
| NPP | X | X | X | X | 234 | Carbon plots | 10 | 10,000 $m^2$ | Subplot | 25 (9 for OP) | 400 $m^2$ | 2011-2016 | No |
| Litterfall | X | X | X | | 8,137 | Carbon plots | 9 | 10,000 $m^2$ | Subplot | 25 | 400 $m^2$ | 2011-2016 | Yes |
| Litter decomposition | X | X | | | 128 | Litter plots | 16 | 6 $m^2$ | Litter bags | 4 | 0.0225 $m^2$ | 2014-2015 | No |

| | | | | | | | | | | | | | |
|---|---|---|---|---|---|---|---|---|---|---|---|---|---|
| Mycelial production | X | X | X | X | 27 | Carbon plots | 9 | 10,000 $m^2$ (3,600 $m^2$ for OP) | Subplot | 3 | 400 $m^2$ | 2016 | No |
| Dung removal | X | X | X | X | 177 | Block (9 2nd-order fractal points) | 15 | ~23.5 ha | Trio of 2nd-order fractal points | 3-13 | ~1.37 ha | 2011-2015 | No |

**Table S2 – Details of Structure & Environment (Level 1) datasets.**

| Dataset | Collector | Description | References for dataset |
|---|---|---|---|
| Above ground carbon | Terhi Riutta | *Data collection* – Above-ground carbon stock (trees >10 cm DBH). All stems (trees >10 cm DBH) within each carbon plot were tagged, identified to species, and measured for diameter and height following the RAINFOR-GEM protocols (*92*). Data were grouped by subplot. Above-ground biomass was estimated using the allometric equation with diameter and height and species-specific wood density by Chave et al. (2005) (*93*). Biomass was converted to carbon by assuming a wood carbon content of 47.4% (*94*).<br><br>*Data preparation* – All data were collected in the carbon plots and were grouped spatially according to Fig. S2. | (*22*) T. Riutta, Y. Malhi, L. K. Kho, T. R. Marthews, W. Huaraca Huasco, M. Khoo, S. Tan, E. Turner, G. Reynolds, S. Both, D. F. R. P. Burslem, Y. A. Teh, C. S. Vairappan, N. Majalap, R. M. Ewers, Logging disturbance shifts net primary productivity and its allocation in Bornean tropical forests. *Global Change Biology*. **24**, 2913–2928 (2018).<br><br>Data are freely available at https://zenodo.org/records/4542881 (*49*). |
| Canopy height | Boris Bongalov, Matheus Nunes, David Milodowski | *Data collection* – Canopy height and vertical profiles of forest structure were compiled using airborne remote sensing with LiDAR collected by NERC's Airborne Research Facility (ARF) in November 2014, using a Leica ALS50-II LiDAR (*95*).<br>A Beer-Lambert approximation was used to convert point clouds to plant area density (PAD) distributions (*95*, *96*), a similar measure to leaf-area index, but where methods do not distinguish between leaves and branches or trunks.<br><br>*Data preparation* – LiDAR measurements for the carbon plots were converted to rasters with 0.5 × 0.5 m cell size. Plots were rotated to a North-South axis if necessary, and then cells were grouped spatially according to Fig. S2. Code and full details for all steps of data manipulation are outlined in the RMarkdown output 'Data_synthesis_analyses/Data preparation/Canopy_height/Processing_canopy_rasters.html' at https://github.com/charliem2003/BALI_synthesis. | (*95*) D. T. Milodowski, D. A. Coomes, T. Swinfield, T. Jucker, T. Riutta, Y. Malhi, M. Svátek, J. Kvasnica, D. F. R. P. Burslem, R. M. Ewers, Y. A. Teh, M. Williams, The impact of logging on vertical canopy structure across a gradient of tropical forest degradation intensity in Borneo. *Journal of Applied Ecology*. **58**, 1764–1775 (2021).<br><br>Data are freely available at https://zenodo.org/records/13329230 (*50*). |
| Leaf-area index | Terhi Riutta | *Data collection* – Leaf area index (LAI) was derived from hemispherical photos (Sigma 8mm SRL fish eye lens and Canon EOS 600D digital camera,<br>mounted on a tripod at 1 m height). Between 5-27 photos were taken over time in each subplot. Images were processed with Hemisfer® software (www.wsl.ch/dienstleistungen/produkte/software/hemisfer/index_EN) (*97*, *98*). LAI was calculated with the method by Thimonier et al. | Data are freely available at https://zenodo.org/records/13329230 (*50*). |

| | | | |
|---|---|---|---|
| | | (2010) (98), with a canopy clumping correction (99) applied.<br><br>*Data preparation* – All data were collected in the carbon plots and were grouped spatially according to Fig. S2. Date was added as a random factor. | |
| Soil temperature | Terhi Riutta | *Data collection* – Soil temperature at 10 cm depth was measured approximately once a month at the centre of each subplot using a hand-held temperature probe (Comark DT400). Data were collected between 3rd Sep. 2011 and 24th Apr. 2017. We used a subset between 26th Mar. 2015 and 3rd Feb. 2017 when all plots were being simultaneously sampled.<br><br>*Data preparation* – All data were collected in the carbon plots and were grouped spatially according to Fig. S2. Date was added as a random factor. | Data are freely available at https://zenodo.org/records/4542881 (49). |
| Soil moisture | Dafydd Elias | *Data collection* – Soil moisture was determined gravimetrically (Dried at 105 °C for 24 hours) from soils sampled in March 2015. Within each plot, five 20×20m subplots were randomly selected. Five soil samples were collected with a 3cm diameter gouge auger within each subplot. The organic soil layer depth was separated from the underlying mineral soil, dried and weighed.<br><br>*Data preparation* – All data were collected in the carbon plots. 5 subplots from each plot were sampled randomly and so Quadrat was not included in the random effects. | Data are freely available at https://catalogue.ceh.ac.uk/documents/7e046092-8405-41b8-9e38-67a844bb9e7d (51). |
| Air temperature: Minimum<br><br>Air temperature: Mean<br><br>Air temperature: Maximum | Benjamin Blonder | *Data collection* – 239 data-loggers recording at synchronised 20 minute intervals for 28 days (start of 1st Nov. 2015 for HLF; 9th Nov. 2015 for MLF and OGF plots). Data-loggers were installed on stakes at the corner and centre of each subplot.<br><br>*Data preparation* – We calculated the daily minimum, maximum and mean temperature for each data-logger. Spatial hierarchy followed the standard clustering of carbon plots (Fig. S2). | (52) B. Blonder, S. Both, D. A. Coomes, D. Elias, T. Jucker, J. Kvasnica, N. Majalap, Y. S. Malhi, D. Milodowski, T. Riutta, M. Svátek, Extreme and highly heterogeneous microclimates in selectively logged tropical forests. *Frontiers in Forests and Global Change*. **1**, 5 (2018).<br><br>Data are freely available in the Supplementary Material of (52). |
| Soil bulk density<br><br>Soil horizon depth<br><br>Soil pH | Dafydd Elias | *Data collection* – Soil sampling was conducted in March 2015. Within each plot, five 20×20m subplots were randomly selected. Five soil samples were collected with a 3cm diameter gouge auger within each subplot. The organic soil layer depth was separated from the underlying mineral soil.<br>Soil pH was measured on fresh soils in a 2.5:1 water: soil slurry suspension, allowed to rest for 30 min and measured using a | Data are freely available at https://catalogue.ceh.ac.uk/documents/7e046092-8405-41b8-9e38-67a844bb9e7d (51). |

| | | |
|---|---|---|
| | | pH meter calibrated between pH 4–7 (pH210 Meter, Hanna Instruments, UK). Total soil C and N was measured using a LECO Truspec Micro elemental analyser (LECO Corporation, USA). Inorganic P was extracted from air-dried soils using a Bray No 1 extractant and analysed using colorimetry on a SEAL AutoAnalyzer 3 (Seal Analytical, UK).<br>An additional soil sample was taken for bulk density using a volumetric ring (7.5cm diameter) with bulk density determined using the volume of the ring and oven-dried soil weight (dried at 105°C for 24 hours) after removal of roots and stones.<br><br>*Data preparation* – All data were collected in the carbon plots. 5 subplots from each plot were sampled randomly and so Quadrat was not included in the random effects. |
| Soil nutrients: C | | |
| Soil nutrients: N | | |
| Soil nutrients: Inorganic P | | |
| Soil nutrients: C:P | | |
| Soil nutrients: C:N | | |

**Table S3 – Details of the tree traits (Level 2).** All tree traits were collected as part of the following study (details in this table have been extracted from table S1 of that publication): S. Both, T. Riutta, C.E.T. Paine, D.M.O. Elias, R.S. Cruz, A. Jain, D. Johnson, U.H. Kritzler, M. Kuntz, N. Majalap-Lee, N. Mielke, M.X. Montoya Pillco, N.J. Ostle, Y. Arn Teh, Y. Malhi, D.F.R.P. Burslem (2019) Logging and soil nutrients independently explain plant trait expression in tropical forests. *New Phytologist*. 221:4, 1853–1865 (*25*).

Methods for generating tree-level trait values and community-weighted mean values for each subplot followed the methods outlined in that publication. 31 physical, chemical and physiological traits were measured on 651 individual trees ≥ 10 cm diameter at breast height (dbh) from 284 tree species. For each trait we generated a community-weighted mean for each sampled subplot by the number of individuals of each species in each subplot and the tree-level trait values. Traits were categorised into three groups by the ecosystem function they most contributed to. For each subplot we also summarised each group as the values from the first principal axis of a principal component analysis (red backgrounds in Fig. S3). . Code and outputs for all steps are outlined in the markdown output 'Data_synthesis_analyses/Data preparation/Tree trait data/Prepare-trait-data.html' at https://zenodo.org/records/13161799 (*47*). Data are freely available at https://zenodo.org/records/3247602 (*53*).

| Trait | Label | Description | Units |
|---|---|---|---|
| **Photosynthetic traits** | | | |
| $\delta^{13}C$ | $\delta^{13}C$ | Indicator of leaf-level water-use efficiency | ‰ |
| Light-saturated photosynthetic rate | Photosynthetic rate: $A_{sat}$ | Measure for metabolic capacity at ambient $CO_2$ concentrations. Proxy for productivity and growth. | µmol CO2 $m^{-2}$ $s^{-1}$ |
| https://zenodo.org/records/3247602Maximum photosynthetic rate | Photosynthetic rate: $A_{max}$ | Measure of the maximum rate at which leaves are able to fix carbon during photosynthesis | µmol CO2 $m^{-2}$ $s^{-1}$ |
| Dark respiration ($R_d$) | Dark respiration | Measure of basal metabolism. Proxy for average realised night-time respiratory carbon flux | µmol CO2 $m^{-2}$ $s^{-1}$ |
| Specific leaf area (SLA) | Specific leaf area | Leaf area divided by oven-dry mass. Proxy for growth and photosynthesis, indicator for carbon investment and leaf longevity | $mm^2$ $mg^{-1}$ |
| $N_a$ | Maximum photosynthetic rate: N (conc) | Close correlation with maximum photosynthetic rate, proxy of nutrient quality for herbivore | mg $mm^{-2}$ |
| $P_a$ | Maximum photosynthetic rate: P (area) | Close correlation with maximum photosynthetic rate, proxy of nutrient quality for herbivore | mg $mm^{-2}$ |
| $Carotenoids_a$ | Carotenoids: Area | Facilitates light-harvesting capacity | mg $mm^{-2}$ |
| $Carotenoids_m$ | Carotenoids: Mass | Facilitates light-harvesting capacity | mg $g^{-1}$ |
| Chlorophyll $a_a$ | Chlorophyll a: Area | Facilitates light-harvesting capacity | mg $mm^{-2}$ |
| Chlorophyll $a_m$ | Chlorophyll a: Mass | Facilitates light-harvesting capacity | mg $g^{-1}$ |
| Chlorophyll $b_a$ | Chlorophyll b: Area | Facilitates light-harvesting capacity | mg $mm^{-2}$ |
| Chlorophyll $b_m$ | Chlorophyll b: Mass | Facilitates light-harvesting capacity | mg $g^{-1}$ |
| **Structural traits** | | | |
| Branch specific density | Branch wood density | Indicator of stability, defence, architecture, hydraulics, carbon gain and growth potential of plants | g $cm^{-3}$ |
| Cellulose | Leaf fibre concentration: Cellulose | Contribution to physical strength and stability of plant cells | % |
| Lignin | Leaf fibre concentration: Lignin | Contribution to physical strength and stability of plant cells | % |
| Hemicellulose | Leaf fibre concentration: Hemicellulose | Contribution to physical strength and stability of plant cells | % |
| Leaf area | Leaf size: Area | Important for light interception and temperature regulation | $mm^2$ |

| | | | |
|---|---|---|---|
| Leaf dry weight | Leaf size: Dry weight | Proxy of leaf size, indicator of water content | mg |
| Leaf force to punch | Leaf strength: Toughness | Physical strength of leaf, indicator for prolonged leaf life span | N $mm^{-1}$ |
| Leaf thickness | Leaf strength: Thickness | Indicator of physical strength of leaf, linked to the number and thickness of mesophyll layers | mm |
| Leaf dry matter content | Leaf strength: Dry matter | Indicator of leaf toughness and leaf lifespan | mg $g^{-1}$ |
| Total phenol concentration | Leaf defence: Phenol | Contribution to plant defence, related to leaf longevity | mg $g^{-1}$ |
| Total tannin concentration | Leaf defence: Tannin | Contribution to plant defence, related to leaf longevity | mg $g^{-1}$ |
| **Nutrients traits** | | | |
| $\delta^{15}N$ | $\delta^{15}N$ | Indicator for nitrogen acquisition via symbiotic fungi and microbes | ‰ |
| $C_m$ | Carbon concentration | Relates to resource capture and defence | % |
| $N_m$ | Maximum photosynthetic rate: N (%) | Close correlation with maximum photosynthetic rate, proxy of nutrient quality for herbivore | % |
| $P_m$ | Maximum photosynthetic rate: P (mass) | Close correlation with maximum photosynthetic rate, proxy of nutrient quality for herbivores | mg $g^{-1}$ |
| $Mg_m$ | Regulatory nutrients: Total Mg | Facilitates function of many cellular enzymes, enables light absorbance in chlorophyll | mg $g^{-1}$ |
| $K_m$ | Regulatory nutrients: Total K | Regulating role for stomata conductance | mg $g^{-1}$ |
| $Ca_m$ | Regulatory nutrients: Total Ca | Regulator for growth processes and responses to environmental stresses, including stomatal function | mg $g^{-1}$ |

**Table S4 – Details of Biodiversity (Level 3) datasets.**

| Dataset | Collector | Description | References for dataset |
|---|---|---|---|
| Soil microbial richness: Protists | Dafydd Elias | *Data collection* – Soil sampling was conducted in March 2015. Within each plot, five 20×20m subplots were randomly selected. Five soil samples were collected with a 3cm diameter gouge auger within each subplot. The organic soil was separated, homogenised, subsampled and transported to the UK Centre for Ecology & Hydrology for microbial analysis.<br>DNA was extracted from 0.2 g soil using a Powersoil® DNA Isolation Kit. Amplicon libraries were constructed according to a dual indexing strategy with each primer consisting of the appropriate Illumina adapter, 8-nt index sequence, a 10-nt pad sequence, a 2-nt linker and the amplicon specific primer (*100*). For bacteria we used V3-V4 16S rRNA amplicon primers following Kozich *et al*. 2013 (*100*). For eukaryotes, the 18S rRNA gene was targeted using amplicon primers following Baldwin *et al.* 2005 (*101*) and fungi were targeted by amplifying the ITS2 region using primers following Ihrmark *et al*. 2012 (*102*).<br>Amplicons were generated using a high fidelity DNA polymerase (Q5 Taq, New England Biolabs). After an initial denaturation at 95°C for 2 minutes, PCR conditions were as follows: denaturation at 95°C for 15 seconds; annealing at temperatures 55°C, 57°C and 52°C for 16S, 18S and ITS reactions respectively; annealing times were 30 seconds with extension at 72°C for 30 seconds; cycle numbers were 25 for 16S and ITS, and 30 for 18S; a final extension of 10 minutes at 72°C was included. Amplicon sizes were determined using an Agilent 2200 TapeStation system and libraries normalized using SequalPrep Normalization Plate Kit (Thermo-Fisher Scientific) and quantified using a Qubit dsDNA HS kit (Thermo-Fisher Scientific). Each amplicon library was sequenced on an Illumina MiSeq using V3 600 cycle reagents at concentrations of 8 pM with a 5% PhiX Illumina control library. Sequencing runs produced in excess of 21, 18 and 16 million reads passing filter for 16S, ITS and 18S amplicons respectively.<br>Sequences were processed in R using DADA2 to quality filter, merge (where appropriate), de-noise and assign taxonomies (*103*). The actual sequence variants (ASV) were subject to taxonomic assignment using the training databases GreenGenes v13.8, Protist Ribosomal Reference database (PR$^2$) v4.12.0 and Unite v7.2 for 16S, 18S and ITS respectively. Fungal taxa were assigned to the ecological guilds ectomycorrhiza, arbuscular mycorrhiza, and all other fungi, using FUNGuild (*104*) . Prior to analysis, samples were normalized by rarefying to 6974, 466 and 10515 reads for 16s, 18s and ITS respectively. | Data are freely available at https://zenodo.org/records/13341608 (*54*). |
| Soil microbial richness: Bacteria | | | |
| Soil fungal richness: Ectomycorrhiza | | | |
| Soil fungal richness: Arbuscular mycorrhiza | | | |

| | | | |
|---|---|---|---|
| Soil richness: Fungi | | *Data preparation* – All data were collected in the carbon plots. Five subplots from each plot were sampled randomly and so we did not include Quadrat in the spatial structure of the random effects in the linear mixed-effects models (see Fig. S2). | |
| Spectral diversity | Matheus Nunes | *Data collection* – Spectral measurements were made on five leaves attached to tree branches used to measure leaf chemical traits. Leaves were randomly selected but we avoided damaged and young plant material to avoid potential confounding factors. Reflectance spectra (350–2500 nm) were acquired using a FieldSpec 4, produced by Analytical Spectral Devices (ASD, Boulder, Colorado, USA). The spectroradiometer's contact probe was mounted on a clamp and firmly pushed down onto the sample against a black background so that no extraneous light was included in the measurement.<br>Spectral measurements were taken halfway between the petiole and leaf tip, and between the main vein and the leaf edge, with the abaxial surface pointing towards the probe. The readings were calibrated against a Spectralon white reference panel every five samples.<br>Leaf reflectance measured at 430 nm, 660 nm, 1450, 1980 nm and 2350 nm align closely with absorption features for pigments, water content, proteins and cellulose (*105*). Spectral diversity calculated from these absorption features can provide an integrated measure of the functional trait variability within plant communities and may be used as a proxy for functional diversity (*105*). In all statistical analyses, the mean reflectance values of the five spectra per branch were used.<br><br>*Data preparation* – For each measurement we calculated the mean across the five spectral bands. We also averaged across multiple measurements for a given branch. All data were collected in the carbon plots and were grouped spatially according to Fig. S2. | Data are freely available at https://zenodo.org/records/13329230 (*50*). |
| Tree diversity: abundance | Sabine Both | *Data collection* – Tree data were collected for all trees ≥ 10 cm diameter breast height (DBH) within carbon plots established as part of the same study used for the tree traits (*25*).<br><br>*Data preparation* – Dead individuals, and those that could not be identified | (*25*) S. Both, T. Riutta, C.E.T. Paine, D.M.O. Elias, R.S. Cruz, A. Jain, D. Johnson, U.H. Kritzler, M. Kuntz, N. Majalap-Lee, N. Mielke, M.X. Montoya |

| | | | |
|---|---|---|---|
| Tree diversity: richness | | to species-level were removed prior to analyses. All data were collected in the carbon plots and were grouped spatially according to Fig. S2.<br><br>*Abundance* – The sum of all individuals ≥ 10 cm DBH within each subplot.<br><br>*Richness* – The total number of species of all individuals ≥ 10 cm DBH within each subplot. | Pillco, N.J. Ostle, Y. Arn Teh, Y. Malhi, D.F.R.P. Burslem. Logging and soil nutrients independently explain plant trait expression in tropical forests. *New Phytologist*. **221**:4, 1853–1865 (2019).<br><br>Data are freely available at https://zenodo.org/records/3247630 (*55*). |
| Liana abundance | Boris Bongalov | *Data collection* – % liana cover for large canopy and emergent trees. The four quadrants of the canopy were scored as 0 (no lianas), 1 (1-20%), 2 (20-40%), 3 (40-60%), 4 (60-80%) and 5 (80-100%).<br><br>*Data preparation* – Scores for the four quadrants per tree were averaged. To allow for logit transformation, trees with scores of 0 were changed to 0.01, and 1 were changed to 0.99. | Data are freely available at https://zenodo.org/records/13329230 (*50*). |
| Dung beetle abundance<br><br>Dung beetle diversity: q = 0<br><br>Dung beetle diversity: q = 1<br><br>Dung beetle diversity: q = 2 | Eleanor Slade | *Data collection* – Sites collected during Feb-March 2011 and 2015 when dung beetle activity peaks using dung-baited pitfall traps (25g human dung for 48 hrs).<br><br>*Data preparation* – Dung beetles were sampled at the 2nd order points of the fractal sampling pattern of the SAFE Project (*14*) allowing us to create three spatial hierarchical levels – Point, Block and Fractal. Points at edge fractals (as opposed to controls) were grouped manually as appropriate.<br>Code and full details for all steps of data manipulation are outlined in the markdown output 'Data_synthesis_analyses/Data preparation/Dung beetle data/Prepare_data.html' at https://zenodo.org/records/13161799 (*47*).<br><br>*Abundance* – Total number of individuals per point across all species.<br><br>*Species richness* – We generated effort-standardised diversity estimates for Hill numbers q=0 (richness), 1 (Shannon) and 2 (Simpsons) using the *iNEXT* package (*74*, *106*) by extrapolating to the mean observed sampling coverage, a more conservative method than that proposed by (*19*). | Data are freely available at https://zenodo.org/record/3247492 (*56*)<br>and<br>https://zenodo.org/record/3247494 (*57*). |
| Bird abundance | Simon Mitchell & Joseph Tobias | *Data collection* – Avian point counts were conducted between 05:50 and 11:00 on days without rain, and were repeated on four separate occasions at each site between 2014 and 2017. During each count, a single experienced observer (SLM / DPE) recorded all bird species heard or seen from the | Data are freely available at https://doi.org/10.5061/dryad.kn251r8 (*58*). |

| | | | |
|---|---|---|---|
| <br>Bird diversity: q = 0<br><br>Bird diversity: q = 1<br><br>Bird diversity: q = 2 | | point for 15 min including fly-overs.<br><br>*Data preparation* – Birds were sampled at the 2nd order points of the fractal sampling pattern of the SAFE Project (*14*) allowing us to create three spatial hierarchical levels – Point, Block and Fractal. Points at edge fractals (as opposed to controls) were grouped manually as appropriate.<br>Code and full details for all steps of data manipulation are outlined in the markdown output 'Data_synthesis_analyses/Data preparation/Bird data/prepare_data.html' at https://zenodo.org/records/13161799 (*47*).<br><br>*Abundance* – Raw total abundance at the point-level.<br><br>*Species richness* – We generated effort-standardised diversity estimates for Hill numbers q=0 (richness), 1 (Shannon) and 2 (Simpsons) using the *iNEXT* package (*74*, *106*) by extrapolating to the mean observed sampling coverage (0.47), a more conservative method than that proposed by (*19*). | |
| Bat abundance<br><br>Bat diversity (small scale)<br><br>Bat diversity (large scale): q = 0<br><br>Bat diversity (large scale): q = 1<br><br>Bat diversity (large scale): q = 2 | David Hemprich-Bennett & Victoria Kemp | *Data collection* – Bats were captured using harp traps placed along flyways (linear features such as paths and streams) (*107*).<br><br>*Data preparation* – Sites were clustered using hierarchical cluster analysis to generate two spatial scales, based on the assumption that ~50% of non-high-flying bat species have home range sizes <25 ha (*108*): (1) 'Group' – using cut-off distance = 250m. Assuming traps sampled a radius of 125m, estimated sampled areas of Groups are 6-16 ha (mean = 9.4 ha); (2) 'Cluster' – estimated sampled areas are 18-28 ha (mean = 23.69 ha).<br>Code for all steps and full details are outlined in the markdown outputs 'Prepare_data_-_species_richness.html', 'Prepare_data_-_beta_diversity.html', 'Prepare_data_-_spatial_grouping.html' and 'home_range_sizes.html' in the folder 'Data_synthesis_analyses/Data preparation/Bat data' at https://zenodo.org/records/13161799 (*47*).<br><br>*Abundance* – Raw total abundance at the point-level (including unidentified species).<br><br>*Species richness* – Unidentified species were first removed. Species richness was calculated at two scales: Small scale: at the point-level (richness only); and Large-scale: at the Group-level. We generated effort- | (*107*) D. R. Hemprich-Bennett, V, A. Kemp, J. Blackman, M. J. Struebig, O. T. Lewis, S. J. Rossiter, E. L. Clare, Altered structure of bat–prey interaction networks in logged tropical forests revealed by metabarcoding. *Molecular Ecology* **30**:22, 5844–57 (2021).<br><br>Data are freely available at https://zenodo.org/records/3247465 (*59*). |

<table>
<tr><td>Bat β-diversity: Nestedness</td><td rowspan="3"></td><td rowspan="3">standardised diversity estimates for Hill numbers q=0 (richness), 1 (Shannon) and 2 (Simpsons) using the iNEXT package (74, 106) by extrapolating to the mean observed sampling coverage (0.84), a more conservative method than that proposed by (19).<br><br>β-diversity – Presence-based Sorensen's dissimilarity was calculated between points within Groups. Total dissimilarity was also partitioned into nestedness and turnover components using the betapart package (66, 109).</td><td rowspan="3"></td></tr>
<tr><td>Bat β-diversity: Turnover</td></tr>
<tr><td>Bat β-diversity: Total</td></tr>
</table>

**Table S5 – Details of Functioning (Level 4) datasets.**

| Dataset | Collector | Description | References for dataset |
|---|---|---|---|
| Respiration: Soil | Terhi Riutta | *Data collection* – $CO_2$ flux from the soil. Soil respiration was measured using a static chamber technique, with one point per subplot. Measurements were made every 4-6 weeks between 2011-2017. Each measurement lasted 124 seconds and the flux was calculated as the slope of a least squares linear regression of the change in $CO_2$ concentration in the chamber headspace over time.<br><br>*Data preparation* – To standardise when data were collected, we retained only subplots with repeated sampling events between 27th Mar. 2015 - 1st Mar. 2017, and Date was used as a random factor. All data were collected in the carbon plots and were grouped spatially according to Fig. S2. | (*110*) T. Riutta, L. K. Kho, Y. A. Teh, R. Ewers, N. Majalap, Y. Malhi, Major and persistent shifts in below-ground carbon dynamics and soil respiration following logging in tropical forests. *Global Change Biology*. **27**, 2225–2240 (2021).<br><br>Data are freely available at https://zenodo.org/records/4542881 (*49*). |
| Respiration: Stem | Terhi Riutta | *Data collection* – Autotrophic respiration of tree trunks was measured every 4-6 weeks using a static chamber method on approximately 40 to 50 stems per plot. Data were collected between 28th Aug. 2011 - 18th Feb. 2017. Each measurement lasted 124 seconds and the flux was calculated as the slope of a least squares linear regression of the change in $CO_2$ concentration in the chamber headspace over time (*93*).<br><br>*Data preparation* – Data collected after 1st Nov. 2016 had potential measurement error and were removed, as well as data after 1st Nov. 2015 for the tower plot, and Date was used as a random factor. All data were collected in the carbon plots and were grouped spatially according to Fig. S2. | Data are freely available at https://zenodo.org/records/12799889 (*60*). |
| NPP | Terhi Riutta | *Data collection* – Data were collected between 2011-2016 over 24 months for each plot and averaged to one observation per subplot.<br><br>*Data preparation* – The NPP measurement here sums canopy, woody and root net primary productivity (the amount of carbon assimilated through photosynthesis that is converted into new tissue, root exudates and volatile organic compounds). All data were collected in the carbon plots and were grouped spatially according to Fig. S2. | (*27*) T. Riutta, Y. Malhi, L. K. Kho, T. R. Marthews, W. Huaraca Huasco, M. Khoo, S. Tan, E. Turner, G. Reynolds, S. Both, D. F. R. P. Burslem, Y. A. Teh, C. S. Vairappan, N. Majalap, R. M. Ewers, Logging disturbance shifts net primary productivity and its allocation in Bornean tropical forests. *Global Change Biology*. **24**, 2913–2928 (2018).<br><br>Data are freely available at https://zenodo.org/records/4542881 (*49*). |
| Litterfall | Terhi Riutta | *Data collection* – Fine litter fall was collected every 14–21 days between 13th Nov. 2011 - 16th May 2016 from 50 × 50 cm litter traps, 1 m above the ground (1 trap per subplot), dried at 70°C until | (*22*) T. Riutta, Y. Malhi, L. K. Kho, T. R. Marthews, W. Huaraca Huasco, M. Khoo, S. Tan, E. Turner, G. Reynolds, S. Both, D. F. R. P. |

| | | | |
|---|---|---|---|
| | | constant weight.<br><br>*Data preparation* – Date was used as a random factor. All data were collected in the carbon plots and were grouped spatially according to Fig. S2. | Burslem, Y. A. Teh, C. S. Vairappan, N. Majalap, R. M. Ewers, Logging disturbance shifts net primary productivity and its allocation in Bornean tropical forests. *Global Change Biology*. **24**, 2913–2928 (2018).<br>Data are freely available at https://zenodo.org/records/4542881 (*49*). |
| Litter decomposition | Sabine Both | *Data collection* – A full-factorial litterbag experiment, with the factors land-use (two levels: OGF and MLF), litter type (two levels: Maliau, SAFE) and mesh size (two levels: fine, coarse). In each land-use type there were 8 pairs of study plots, consisting of one plot per litter type each. 2 × 3 m plots were established in Oct. 2014, protected against litter fall by suspending a fishing net 50 cm above.<br><br>*Data preparation* – The factorial design was used as the spatial hierarchy for the random effects structure –<br>(1 \| Plot / Litter type / Mesh size). | (*111*) S. Both, D. M. O. Elias, U. H. Kritzler, N. J. Ostle, D. Johnson, Land use not litter quality is a stronger driver of decomposition in hyperdiverse tropical forest. *Ecology and Evolution*. **7**, 9307–9318 (2017).<br><br>Data are freely available at https://zenodo.org/records/3247638 (*61*). |
| Mycelial production | Samuel Robinson | *Data collection* – Three 20 × 20 m subplots were randomly chosen per 1 ha plot. Ten hyphal in-growth bags were installed per subplot at a randomly chosen location. Bags were buried at 50 cm intervals along two parallel transects (five bags per transect) spaced 1 m apart. In-growth bags were made with 41 µm nylon mesh filled with sterilised quartz sand and left in place for 6 months. Where possible, sand from three undamaged bags in each subplot were bulked and homogenised for analysis so that there remains one measurement per subplot.<br><br>*Data preparation* – As the three sampled subplots from each plot were homogenised we did not include Quadrat or Subplot in the spatial structure of the random effects in the linear mixed-effects models. | (*112*) S. J. B. Robinson, D. Elias, D. Johnson, S. Both, T. Riutta, T. Goodall, N. Majalap, N. P. McNamara, R. Griffiths, N. Ostle, Soil fungal community characteristics and mycelial production across a disturbance gradient in lowland Dipterocarp rainforest in Borneo. *Frontiers in Forests and Global Change* **3**:64 (2020).<br><br>Data are freely available at 10.5281/zenodo.13122107 (*62*). |
| Dung removal | Eleanor Slade | *Data collection* – Sites were sampled in Feb-March 2011 and 2015 when dung beetle activity peaks (single year-sampled sites were removed). 700 g of cattle dung was left for 24 hours and the mass of dung removed measured (minus mass loss due to evaporation measured from beetle-excluded dung) (*35*).<br><br>*Data preparation* – Dung beetles were sampled at the 2$^{nd}$ order points of the fractal sampling pattern of the SAFE Project (*14*) allowing us to create three spatial hierarchical levels – Point, Block and Fractal. Points at edge fractals (as opposed to controls) were grouped manually as appropriate. | Data are freely available at https://zenodo.org/record/3247492 (*56*) and https://zenodo.org/record/3247494 (*57*). |

**Table S6 – Outputs from the best model for each dataset**. For each disturbance category (OGF = old-growth forest; MLF = moderately logged forest; HLF = highly logged forest; OP = oil palm plantation) the mean effects sizes ± 95 % confidence intervals, along with the *p*-value and marginal $R^2$ of the overall model are given. The full summary output for each model can be seen in the RMarkdown documents accompanying each dataset, available at https://zenodo.org/records/13161799 (*47*).

| **Dataset** | ***p*** | **Marg. $R^2$** | **OGF** | **MLF** | **HLF** | **OP** |
|---|---|---|---|---|---|---|
| ***Structure & Environment*** | | | | | | |
| Above ground carbon | <0.001 | 0.395 | 0.755 (0.557 - 0.954) | 0.153 (-0.376 - 0.683) | -0.975 (-1.494 - -0.456) | -1.136 (-1.789 - -0.484) |
| Canopy height | <0.001 | 0.358 | 0.547 (0.302 - 0.792) | -0.554 (-1.224 - 0.115) | -1.056 (-1.725 - -0.387) | |
| Leaf-area Index | <0.001 | 0.177 | 0.467 (0.291 - 0.643) | 0.161 (-0.335 - 0.657) | -0.407 (-0.828 - 0.015) | -0.911 (-1.554 - -0.268) |
| Soil temperature | 0.413 | 0.037 | -0.55 (-1.073 - -0.028) | -0.329 (-1.778 - 1.121) | -0.12 (-1.43 - 1.19) | 0.183 (-1.504 - 1.869) |
| Soil moisture | 0.044 | 0.188 | -0.093 (-0.663 - 0.476) | 0.237 (-1.347 - 1.822) | 0.528 (-1.033 - 2.089) | -1.312 (-3.151 - 0.526) |
| Air temperature: Min. | <0.001 | 0.33 | 0.295 (0.075 - 0.515) | 0.447 (0.008 - 0.885) | -0.852 (-1.293 - -0.411) | |
| Air temperature: Mean | <0.001 | 0.146 | -0.156 (-0.418 - 0.107) | 0.475 (-0.044 - 0.994) | -0.421 (-0.943 - 0.101) | |
| Air temperature: Max. | 0.002 | 0.08 | -0.417 (-0.687 - -0.147) | 0.299 (-0.324 - 0.923) | 0.077 (-0.551 - 0.706) | |
| Soil bulk density | <0.001 | 0.294 | -0.665 (-1.016 - -0.314) | 0.622 (-0.317 - 1.56) | 0.231 (-0.7 - 1.163) | 1.085 (-0.037 - 2.207) |
| Soil horizon depth | <0.001 | 0.25 | -0.422 (-0.678 - -0.166) | 0.74 (0.001 - 1.479) | 0.482 (-0.282 - 1.247) | -0.858 (-1.738 - 0.022) |
| Soil pH | 0.044 | 0.27 | 0.358 (-0.322 - 1.038) | -1.144 (-2.993 - 0.705) | 0.188 (-1.665 - 2.041) | 0.479 (-1.711 - 2.669) |
| Soil nutrients: C | 0.487 | 0.06 | -0.157 (-0.788 - 0.474) | 0.441 (-1.312 - 2.194) | 0.075 (-1.648 - 1.799) | -0.405 (-2.479 - 1.669) |
| Soil nutrients: N | 0.808 | 0.025 | -0.193 (-0.871 - 0.484) | 0.188 (-1.681 - 2.056) | 0.139 (-1.712 - 1.989) | 0.121 (-2.095 - 2.338) |
| Soil nutrients: C:N | 0.03 | 0.214 | 0.02 (-0.513 - 0.554) | 0.697 (-0.774 - 2.167) | -0.109 (-1.561 - 1.343) | -1.255 (-2.994 - 0.483) |
| Soil nutrients: Inorganic P | 0.029 | 0.265 | -0.416 (-0.969 - 0.137) | 0.458 (-1.055 - 1.971) | -0.263 (-1.771 - 1.246) | 1.272 (-0.55 - 3.095) |
| Soil nutrients: C:P | 0.008 | 0.339 | 0.363 (-0.163 - 0.889) | -0.187 (-1.618 - 1.243) | 0.364 (-1.065 - 1.793) | -1.805 (-3.541 - -0.069) |
| ***Photosynthesis traits*** | | | | | | |
| Photosyn. traits | 0.002 | 0.289 | -0.596 (-0.991 - -0.202) | 0.319 (-0.771 - 1.409) | 0.883 (-0.227 - 1.993) | |
| δ13C | 0.002 | 0.298 | -0.552 (-0.955 - -0.149) | 0.079 (-1.057 - 1.215) | 1.037 (-0.093 - 2.167) | |
| Photosyn. rate: Asat | 0.006 | 0.29 | -0.614 (-1.084 - -0.144) | 0.359 (-0.955 - 1.672) | 0.877 (-0.45 - 2.203) | |

| Dataset | *p* | Marg. $R^2$ | OGF | MLF | HLF | OP |
|---|---|---|---|---|---|---|
| Photosyn. rate: Amax | 0.006 | 0.279 | -0.596 (-1.066 - -0.126) | 0.313 (-1.001 - 1.627) | 0.884 (-0.439 - 2.207) | |
| Max. photosyn. rate: N(conc) | 0.029 | 0.175 | -0.331 (-0.825 - 0.162) | -0.145 (-1.515 - 1.226) | 0.817 (-0.532 - 2.166) | |
| Max. photosyn. rate: P(area) | 0.076 | 0.143 | 0.10 (-0.442 - 0.659) | -0.683 (-2.19 - 0.825) | 0.466 (-1.029 - 1.962) | |
| Dark respiration | 0.006 | 0.227 | 0.424 (0.035 - 0.812) | 0.041 (-1.05 - 1.131) | -0.889 (-1.992 - 0.214) | |
| Specific leaf area | 0.541 | 0.016 | 0.134 (-0.27 - 0.538) | -0.135 (-1.227 - 0.957) | -0.122 (-1.224 - 0.98) | |
| Carotenoids: Area | 0.298 | 0.015 | 0.105 (-0.088 - 0.299) | -0.194 (-0.811 - 0.423) | -0.018 (-0.582 - 0.547) | |
| Carotenoids: Mass | 0.21 | 0.034 | 0.171 (-0.164 - 0.507) | -0.285 (-1.206 - 0.635) | -0.052 (-0.97 - 0.866) | |
| Chlorophyll a: Area | 0.024 | 0.059 | 0.237 (0.015 - 0.458) | -0.348 (-1.034 - 0.337) | -0.129 (-0.75 - 0.491) | |
| Chlorophyll a: Mass | 0.025 | 0.087 | 0.294 (-0.017 - 0.604) | -0.431 (-1.282 - 0.42) | -0.155 (-0.995 - 0.686) | |
| Chlorophyll b: Area | <0.001 | 0.227 | 0.531 (0.314 - 0.748) | -0.421 (-1.062 - 0.219) | -0.651 (-1.277 - -0.025) | |
| Chlorophyll b: Mass | 0.002 | 0.212 | 0.513 (0.184 - 0.842) | -0.449 (-1.348 - 0.451) | -0.581 (-1.487 - 0.326) | |
| ***Nutrient traits*** | | | | | | |
| Nutrient traits | 0.027 | 0.197 | 0.007 (-0.479 - 0.492) | -0.693 (-2.02 - 0.635) | 0.68 (-0.671 - 2.032) | |
| δ15N | 0.024 | 0.181 | 0.468 (-0.001 - 0.937) | -0.342 (-1.678 - 0.994) | -0.598 (-1.888 - 0.692) | |
| Carbon conc. | 0.078 | 0.136 | -0.124 (-0.648 - 0.399) | 0.659 (-0.775 - 2.093) | -0.434 (-1.894 - 1.026) | |
| Max. photosyn. rate: N(%) | 0.008 | 0.205 | -0.324 (-0.712 - 0.065) | -0.199 (-1.264 - 0.866) | 0.868 (-0.229 - 1.965) | |
| Max. photosyn. rate: P(mass) | 0.09 | 0.157 | 0.154 (-0.464 - 0.772) | -0.744 (-2.424 - 0.936) | 0.445 (-1.238 - 2.129) | |
| Regulat. nutrients: Total Mg | 0.369 | 0.053 | 0.137 (-0.434 - 0.707) | -0.423 (-1.98 - 1.133) | 0.154 (-1.419 - 1.726) | |
| Regulat. nutrients: Total K | 0.02 | 0.137 | -0.205 (-0.598 - 0.188) | -0.269 (-1.337 - 0.8) | 0.699 (-0.367 - 1.764) | |
| Regulat. nutrients: Total Ca | 0.146 | 0.132 | 0.232 (-0.419 - 0.883) | -0.7 (-2.468 - 1.067) | 0.255 (-1.542 - 2.052) | |
| ***Structural traits*** | | | | | | |
| Structural traits | 0.002 | 0.274 | 0.484 (0.115 - 0.852) | -0.036 (-1.062 - 0.991) | -0.945 (-1.965 - 0.076) | |
| Branch wood density | <0.001 | 0.392 | 0.654 (0.352 - 0.957) | -0.161 (-0.994 - 0.673) | -1.169 (-1.995 - -0.344) | |

| Dataset | *p* | Marg. $R^2$ | OGF | MLF | HLF | OP |
|---|---|---|---|---|---|---|
| Leaf fibre conc.: Cellul. | 0.052 | 0.175 | 0.444<br>(-0.113 - 1) | -0.234<br>(-1.765 - 1.296) | -0.662<br>(-2.183 - 0.86) | |
| Leaf fibre conc.: Lignin | 0.061 | 0.165 | 0.366<br>(-0.186 - 0.917) | 0.01<br>(-1.501 - 1.521) | -0.743<br>(-2.258 - 0.772) | |
| Leaf fibre conc.: Hemicel. | 0.198 | 0.096 | 0.328<br>(-0.262 - 0.918) | -0.245<br>(-1.859 - 1.368) | -0.426<br>(-2.045 - 1.193) | |
| Leaf size: Area | 0.107 | 0.07 | -0.139<br>(-0.525 - 0.246) | -0.197<br>(-1.292 - 0.898) | 0.483<br>(-0.59 - 1.555) | |
| Leaf size: Dry wgt | 0.108 | 0.066 | -0.148<br>(-0.527 - 0.231) | -0.171<br>(-1.249 - 0.907) | 0.471<br>(-0.583 - 1.525) | |
| Leaf strength: Tough. | 0.01 | 0.222 | 0.526<br>(0.08 - 0.972) | -0.449<br>(-1.674 - 0.777) | -0.616<br>(-1.836 - 0.605) | |
| Leaf strength: Thick. | 0.031 | 0.122 | -0.342<br>(-0.727 - 0.043) | 0.132<br>(-0.947 - 1.211) | 0.56<br>(-0.51 - 1.629) | |
| Leaf strength: Dry mat. | 0.12 | 0.074 | 0.107<br>(-0.305 - 0.519) | 0.253<br>(-0.878 - 1.385) | -0.49<br>(-1.631 - 0.65) | |
| Leaf defence: Phenol | 0.007 | 0.213 | -0.509<br>(-0.91 - -0.109) | 0.679<br>(-0.452 - 1.81) | 0.34<br>(-0.774 - 1.454) | |
| Leaf defence: Tannin | 0.046 | 0.074 | -0.229<br>(-0.563 - 0.106) | 0.461<br>(-0.433 - 1.356) | -0.003<br>(-0.929 - 0.922) | |
| ***Biodiversity*** | | | | | | |
| Soil microbial richness: Bacteria | <0.001 | 0.291 | 0.175<br>(-0.16 - 0.511) | -1.125<br>(-2.028 - -0.223) | 0.392<br>(-0.526 - 1.311) | 0.715<br>(-0.412 - 1.842) |
| Soil microbial richness: Protist | 0.146 | 0.089 | 0.232<br>(-0.251 - 0.715) | -0.533<br>(-1.85 - 0.785) | 0.197<br>(-1.109 - 1.503) | -0.302<br>(-1.828 - 1.225) |
| Soil fungal richness.: Ectomycorrhiza | <0.001 | 0.362 | 0.758<br>(0.593 - 0.923) | -0.529<br>(-1.064 - 0.005) | -0.323<br>(-0.784 - 0.137) | -1.282<br>(-1.882 - -0.682) |
| Soil fungal rich.: Arbusc. mycorrhiza | 0.063 | 0.09 | -0.101<br>(-0.488 - 0.286) | -0.313<br>(-1.367 - 0.741) | 0.574<br>(-0.472 - 1.619) | -0.158<br>(-1.395 - 1.08) |
| Soil fungal richness: Fungi | 0.002 | 0.154 | 0.141<br>(-0.126 - 0.408) | -0.617<br>(-1.369 - 0.136) | 0.547<br>(-0.171 - 1.266) | -0.444<br>(-1.278 - 0.39) |
| Spectral diversity | 0.216 | 0.037 | -0.088<br>(-0.427 - 0.252) | -0.453<br>(-1.398 - 0.492) | 0.076<br>(-0.851 - 1.004) | |
| Tree diversity: abundance | 0.033 | 0.079 | -0.014<br>(-0.274 - 0.245) | 0.415<br>(-0.331 - 1.162) | -0.386<br>(-1.156 - 0.383) | |
| Tree diversity: richness | <0.001 | 0.140 | 0.053<br>(-0.137 - 0.244) | 0.498<br>(-0.010 - 1.096) | -0.605<br>(-1.192 - -0.018) | |
| Liana abundance | 0.032 | 0.079 | -0.678<br>(-1.345 - -0.01) | 0.245<br>(-1.198 - 1.688) | 0.017<br>(-1.441 - 1.476) | |
| Dung beetle abundance | <0.001 | 0.319 | 0.728<br>(0.315 - 1.142) | -0.124<br>(-1.038 - 0.79) | 0.252<br>(-0.614 - 1.118) | -1.483<br>(-2.354 - -0.613) |
| Dung beetle diversity: $q$=0 | <0.001 | 0.279 | 0.597<br>(0.288 - 0.907) | 0.729<br>(-0.014 - 1.472) | -0.019<br>(-0.703 - 0.664) | -1.267<br>(-2.05 - -0.484) |
| Dung beetle diversity: $q$=1 | <0.001 | 0.25 | 0.623<br>(0.173 - 1.073) | 0.742<br>(-0.325 - 1.808) | -0.086<br>(-1.047 - 0.876) | -1.071<br>(-2.077 - -0.066) |

| Dataset | *p* | Marg. $R^2$ | OGF | MLF | HLF | OP |
|---|---|---|---|---|---|---|
| Dung beetle diversity: *q*=2 | <0.001 | 0.216 | 0.613 (0.118 - 1.108) | 0.576 (-0.626 - 1.778) | -0.053 (-1.116 - 1.009) | -1.01 (-2.124 - 0.103) |
| Bird abund. | <0.001 | 0.166 | 0.361 (-0.02 - 0.743) | -0.125 (-1.024 - 0.775) | 0.11 (-0.719 - 0.938) | -1.226 (-2.181 - -0.271) |
| Bird diversity: *q*=0 | <0.001 | 0.266 | 0.423 (0.05 - 0.797) | 0.545 (-0.31 - 1.4) | -0.046 (-0.827 - 0.735) | -1.598 (-2.443 - -0.753) |
| Bird diversity: *q*=1 | <0.001 | 0.265 | 0.408 (0.038 - 0.777) | 0.567 (-0.277 - 1.41) | -0.044 (-0.814 - 0.726) | -1.596 (-2.426 - -0.766) |
| Bird diversity: *q*=2 | <0.001 | 0.264 | 0.338 (-0.028 - 0.704) | 0.591 (-0.242 - 1.424) | -0.014 (-0.78 - 0.752) | -1.616 (-2.443 - -0.79) |
| Bat abund. | 0.002 | 0.053 | 0.364 (0.15 - 0.577) | -0.151 (-0.722 - 0.421) | -0.159 (-0.635 - 0.316) | |
| Bat diversity (sm scale) | 0.013 | 0.027 | 0.229 (0.039 - 0.42) | 0.086 (-0.439 - 0.611) | -0.14 (-0.566 - 0.285) | |
| Bat diversity (lg scale): *q*=0 | 0.024 | 0.192 | -0.312 (-0.873 - 0.25) | 1.206 (-0.483 - 2.896) | -0.117 (-1.379 - 1.146) | |
| Bat diversity (lg scale): *q*=1 | 0.007 | 0.249 | -0.415 (-0.818 - -0.012) | 1.383 (-0.107 - 2.873) | -0.077 (-1.006 - 0.852) | |
| Bat diversity (lg scale): *q*=2 | 0.005 | 0.223 | -0.436 (-0.821 - -0.051) | 1.262 (-0.058 - 2.583) | -0.017 (-0.91 - 0.876) | |
| Bat β-diversity: Nestedness | 0.019 | 0.09 | 0.378 (-0.143 - 0.898) | -0.571 (-1.702 - 0.56) | -0.11 (-1.248 - 1.028) | |
| Bat β-diversity: Turnover | 0.019 | 0.092 | -0.37 (-0.903 - 0.163) | 0.606 (-0.557 - 1.77) | 0.096 (-1.06 - 1.253) | |
| Bat β-diversity: Total | 0.047 | 0.095 | -0.386 (-0.77 - -0.002) | -0.192 (-1.179 - 0.795) | 0.298 (-0.648 - 1.243) | |
| ***Functioning*** | | | | | | |
| Respiration: Soil | 0.005 | 0.036 | -0.059 (-0.232 - 0.114) | 0.17 (-0.315 - 0.667) | 0.029 (-0.384 - 0.441) | -0.716 (-1.32 - -0.111) |
| Respiration: Stem | <0.001 | 0.042 | -0.398 (-0.577 - -0.219) | 0.142 (-0.307 - 0.59) | 0.049 (-0.37 - 0.467) | |
| NPP | 0.090 | 0.164 | -0.119 (-0.453 - 0.215) | 0.435 (-0.494 - 1.365) | -0.195 (-1.101 - 0.712) | 0.518 (-0.683 - 1.720) |
| Litterfall | 0.003 | 0.097 | 0.465 (0.206 - 0.723) | 0.01 (-0.681 - 0.702) | -0.31 (-0.952 - 0.332) | |
| Litter decomp. | <0.001 | 0.121 | 0.364 (0.064 - 0.664) | -0.36 (-1.053 - 0.324) | | |
| Mycelial production | 0.012 | 0.133 | 0.164 (-0.371 - 0.698) | 0.282 (-1.537 - 2.102) | -0.098 (-1.32 - 1.124) | -1.023 (-2.125 - 0.079) |
| Dung removal | 0.161 | 0.089 | 0.696 (-0.001 - 1.393) | 0.038 (-1.642 - 1.717) | -0.217 (-1.757 - 1.322) | 0.038 (-1.622 - 1.699) |
